# Post-transcriptional gene regulation by the RNA binding protein IGF2BP3 is critical for MLL-AF4 mediated leukemogenesis

**DOI:** 10.1101/2020.12.20.423624

**Authors:** Tiffany M Tran, Julia Philipp, Jaspal Bassi, Neha Nibber, Jolene Draper, Tasha Lin, Jayanth Kumar Palanichamy, Amit Kumar Jaiswal, Oscar Silva, May Paing, Jennifer King, Sol Katzman, Jeremy R. Sanford, Dinesh S. Rao

**Affiliations:** Department of Pathology and Laboratory Medicine, David Geffen School of Medicine at UCLA, Los Angeles, California, USA; Molecular, Cellular and Integrative Physiology Graduate Program, UCLA, Los Angeles, California, 90095, USA; Department of Molecular, Cellular and Developmental Biology, UCSC, Santa Cruz, California, 95064, USA; Division of Hematology/Oncology, Department of Medicine at UCLA, Los Angeles, California, 90095, USA; Molecular Biology Graduate Program, UCLA, Los Angeles, California, 90095, USA; Department of Biochemistry, All India Institute of Medical Sciences, New Delhi, 110029, India; Department of Pathology, Stanford University School of Medicine, Stanford, California, 94305, USA; Division of Rheumatology, Department of Medicine at UCLA, Los Angeles, California, 90095, USA; Jonsson Comprehensive Cancer Center (JCCC); Broad Stem Cell Research Center, UCLA, Los Angeles, California, 90095, USA

## Abstract

Despite recent advances in therapeutic approaches, patients with MLL-rearranged leukemia still have poor outcomes and a high risk of relapse. Here, we found that MLL-AF4, the most common MLL fusion protein in patients, transcriptionally induces IGF2BP3 and that IGF2BP3 strongly amplifies MLL-Af4 mediated leukemogenesis. Deletion of *Igf2bp3* significantly increases the survival of mice with MLL-Af4 driven leukemia and greatly attenuates disease, with a minimal impact on baseline hematopoiesis. At the cellular level, MLL-Af4 leukemia-initiating cells require *Igf2bp3* for their function in leukemogenesis. eCLIP and transcriptome analysis of MLL-Af4 transformed stem and progenitor cells and MLL-Af4 bulk leukemia cells reveals a complex IGF2BP3-regulated post-transcriptional operon governing leukemia cell survival and proliferation. Regulated mRNA targets include important leukemogenic genes such as those in the *Hoxa* locus and numerous genes within the Ras signaling pathway. Together, our findings show that IGF2BP3 is an essential positive regulator of MLL-AF4 mediated leukemogenesis and represents an attractive therapeutic target in this disease.

## INTRODUCTION

Chromosomal rearrangements of the mixed-lineage leukemia (*MLL*, also known as *KMT2A*) gene are recurrently found in a subset of acute lymphoblastic leukemia (ALL), acute myeloid leukemia (AML), and in acute leukemia of ambiguous lineage (Krivtsov and Armstrong, 2007). Despite recent advances in therapeutic approaches, patients with *MLL*-rearranged (MLLr) leukemia have very poor outcomes, a high risk of relapse, and show resistance to novel targeted therapies (Moorman et al., 2010; Pui et al., 2011) (Haddox et al., 2017; Rayes et al., 2016; Wei et al., 2008). *MLL* encodes a H3K4 methyltransferase that has been shown to be required for hematopoietic stem cell (HSC) development during both embryonic and adult hematopoiesis (Ernst et al., 2004a; Jude et al., 2007). Many of the translocation partners for *MLL*, including *AF4 (AFF1)*, encode proteins that regulate transcriptional elongation (Lin et al., 2010; Mohan et al., 2010; Smith et al., 2011). Of more than 90 translocation fusion partner genes, *MLL-AF4* (*KMT2A-AFF1*) is the most common *MLL* fusion protein in patients (Meyer et al., 2018). Biologically, *MLL*-*AF4*-driven leukemia is a distinct entity, with a unique gene expression profile showing significant overlap with stem cell programs (Krivtsov et al., 2008; Somervaille et al., 2009).

At the post-transcriptional level, emerging evidence suggests a role for microRNAs, RNA-binding proteins, and other novel mechanisms in regulating gene expression during leukemogenesis (Ennajdaoui et al., 2016; Jønson et al., 2014; Nguyen et al., 2014; Palanichamy et al., 2016; Park et al., 2015). We recently identified the oncofetal RNA binding protein (RBP) Insulin like growth factor 2 mRNA binding protein 3 (IGF2BP3) as an important regulator of gene expression in *MLL*-rearranged B-ALL (Palanichamy et al., 2016). IGF2BP3 is expressed during embryogenesis, lowly expressed in healthy adult tissues, and strongly re-expressed in cancer cells (Mueller et al., 2003; Mueller-Pillasch et al., 1999). Several studies have shown that elevated levels of IGF2BP3 expression are correlated with diminished patient survival in many cancer types and may be a marker of disease aggressiveness in B-ALL (Kobel et al., 2009; Lochhead et al., 2012; Schaeffer et al., 2010; Stoskus et al., 2011). Previously, we determined that overexpression of IGF2BP3 in the bone marrow (BM) of mice leads to a pathologic expansion of hematopoietic stem and progenitor cells (HSPC), in a manner dependent on RNA binding. IGF2BP3 interacts primarily with the 3’UTR of its target transcripts, as with MYC and CDK6, resulting in an upregulation of transcript and protein (Palanichamy et al., 2016). In the case of MYC and CDK6, IGF2BP3 binding resulted in upregulation of target transcript and protein, with attendant effects on pathologic hematopoietic stem and progenitor cell expansion. Together, these studies illuminated a novel role for post-transcriptional gene regulation in the pathologic proliferation of HSPCs.

Experimentally, MLL-AF4 driven leukemogenesis has been studied using a range of *in vitro* and *in vivo* models leading to significant progress in our understanding of MLL-rearranged leukemia (Bursen et al., 2010; Chen et al., 2006; Krivtsov et al., 2008; Metzler et al., 2006; Montes et al., 2011; Tamai et al., 2011). Here, we explicitly tested the requirement for *Igf2bp3* in a bona-fide *in vivo* model of MLL-Af4 driven leukemogenesis (Lin et al., 2016). Deletion of *Igf2bp3* significantly increased the survival of MLL-Af4 transplanted mice and decreased the numbers and self-renewal capacity of MLL-Af4 leukemia-initiating cells (LICs). Mechanistically, we found that IGF2BP3 targets and modulates the expression of transcripts within the *Hoxa* locus and components of the Ras signaling pathway, both key regulators of leukemogenesis, through multiple post-transcriptional mechanisms (Downward, 2003; Milne et al., 2010). Together, our findings show that IGF2BP3 is a critical regulator of MLL-AF4 mediated leukemogenesis and a potential therapeutic target in this disease (Downward, 2003; Milne et al., 2010).

## RESULTS

### The MLL-AF4 fusion protein transcriptionally induces IGF2BP3

To determine the functional impact of IGF2BP3 expression on MLL-AF4-mediated gene expression, we compared IGF2BP3-regulated targets with a published dataset of MLL-Af4 targets obtained by ChIP-Seq (Lin et al., 2016; Palanichamy et al., 2016). Transcripts modulated by IGF2BP3 were significantly enriched for MLL-Af4-bound genes (Figure 1A; Supplemental Figure 1A). Interestingly, IGF2BP3 itself was a direct transcriptional target of MLL-Af4, with binding sites within the first intron and promoter region of IGF2BP3 (Lin et al., 2016). To confirm if *IGF2BP3* was a direct transcriptional target of MLL-AF4, we performed ChIP-PCR assays on RS4;11 and SEM cell lines, human B-ALL cell lines that contain the MLL-AF4 translocation, and determined that a region in the first intron of *IGF2BP3* is strongly bound by MLL-AF4 (Figure 1B; Supplemental Figure 1B) (Wilkinson et al., 2013). This MLL-AF4 binding was abrogated when SEM cells were treated with the bromodomain inhibitor, iBET-151 (Supplemental Figure 1C) (Dawson et al., 2011). Furthermore, we observed an MLL-AF4-dose-dependent increase in luciferase reporter activity, using a 950bp promoter region upstream of the transcription start site (TSS) of IGF2BP3 (Figure 1C). To confirm that MLL-AF4 not only binds to the *IGF2BP3* gene but also promotes its expression, we utilized a retroviral MSCV vector encoding the human *MLL* fused to the murine *Af4* (MLL-Af4)(Lin et al., 2016). In the murine pre-B cell line, 70Z/3, and primary murine bone marrow cells, we found that MLL-Af4 transduction caused an approximately 64-fold upregulation of *Igf2bp3* mRNA (Figure 1D-E). Concordantly, IGF2BP3 protein was upregulated in MLL-Af4 transduced primary bone marrow cells, after being undetectable in control cells (Figure 1F). Taken together, these findings demonstrate that MLL-Af4 drives the expression of *Igf2bp3 in vivo*.

**Figure 1:**
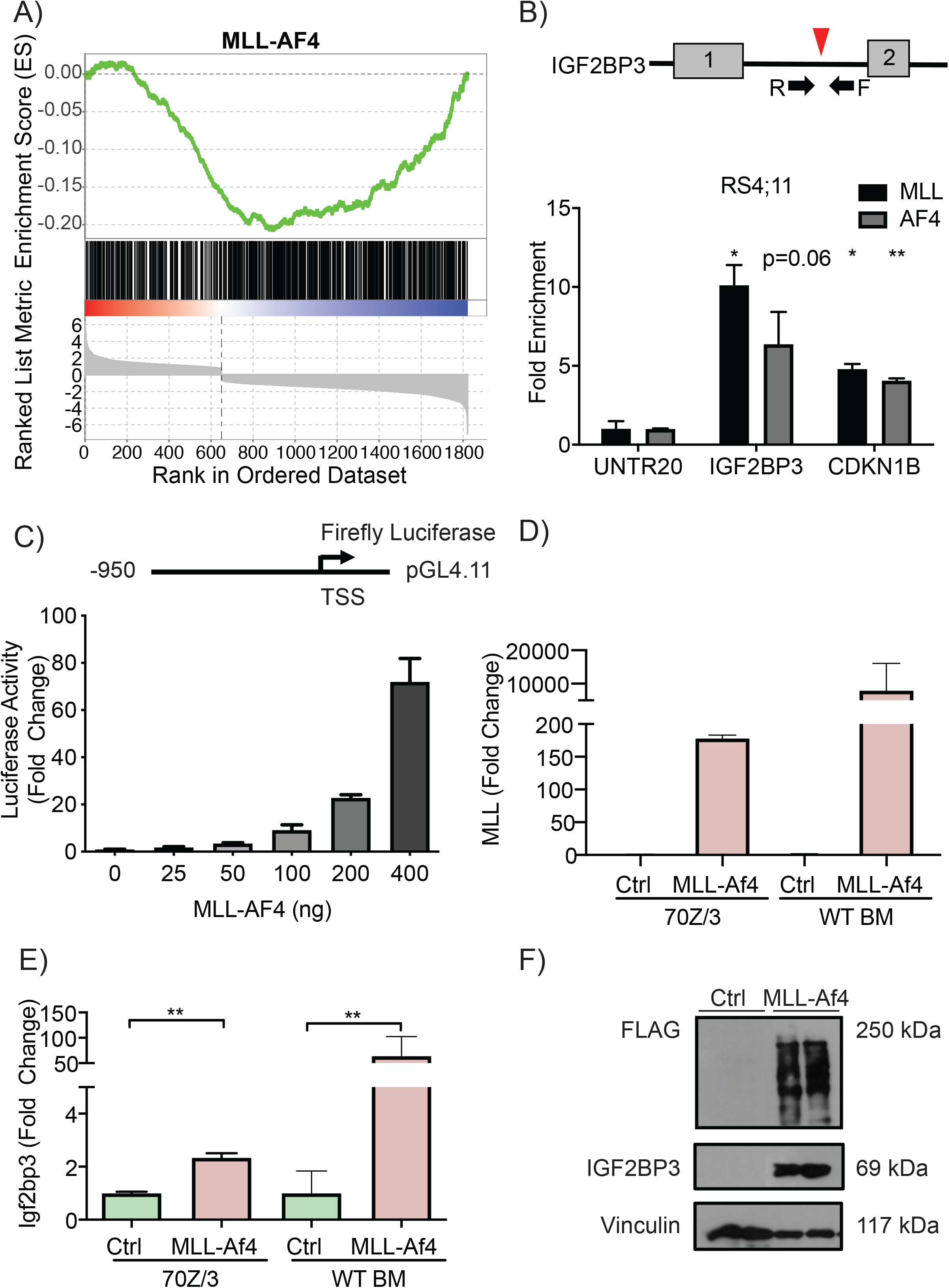
MLL-AF4 transcriptionally induces IGF2BP3. A) GSEA of differentially expressed genes from IGF2BP3 depleted RS4;11 cells shows significant negative enrichment with MLL-AF4 ChIP targets (nominal P value: 0.001, FDR: 0.001, Normalized ES: -1.54)). B) Schematic of MLL-AF4 binding site in intron 1 of IGF2BP3 (top). ChIP-qPCR shows fold enrichment for IGF2BP3 and CDKN1B with MLL and AF4 IP in RS4;11. Normalized to UNTR20, an untranscribed region (t-test; *P < 0.05, **P < 0.01). C) Luciferase assay of the IGF2BP3 promoter shows a dose-dependent response to MLL-AF4. D) Expression of MLL through RT-qPCR of 70Z/3 transduced with either control (Ctrl) or MLL-Af4 vector selected with G418 and MLL expression at the RNA level in the BM of WT recipients transplanted with Ctrl or MLL-Af4 HSPCs. E) Induction of Igf2bp3 at the RNA level in selected 70Z/3 with MLL-Af4 and in the BM of WT recipients transplanted with Ctrl or MLL-Af4 HSPCs (bottom) (t-test; **P < 0.01). F) Induction of Igf2bp3 at the protein level in BM from mice transplanted with MLL-Af4 transduced WT donor HSPCs.

### Normal hematopoiesis is maintained in Igf2bp3 KO mice

To test the *in vivo* requirement for IGF2BP3 in leukemogenesis, we generated an *Igf2bp3* KO (I3KO) mouse. We initially generated a floxed *Igf2bp3* allele (f/f; Supplemental Figure 2A) using CRISPR/Cas9. In the course of mating the mice with the Vav1-Cre mouse strain, we serendipitously generated a germline knockout allele (del), which we isolated and further characterized (Supplemental Figure 2B). This generation of a germline knockout allele is consistent with prior reports for the Vav1-Cre mouse strain, which displays “leaky” Cre expression resulting in germline deletion (Croker et al., 2004; de Boer et al., 2003; Georgiades et al., 2002; Heffner et al., 2012; Joseph et al., 2013). Mendelian inheritance was confirmed, although the distribution of genotypes was marginally skewed (Table S1). Deletion of *Igf2bp3* was confirmed at the DNA, RNA, and protein level (Supplemental Figure 2C-E). Thus, *Igf2bp3*^*del/del*^ (I3KO) mice were used for the remainder of the study. Immunophenotyping of I3KO mice showed no significant differences in the numbers of HSPCs in the BM compared to WT (Supplemental Figure 2F). Additionally, I3KO mice showed similar numbers of myeloid-lineage progenitors (including CMPs, GMPs, and MEPs)(Supplemental Figure 2G) and normal B-cell development as assessed by the Hardy scheme (Hardy and Hayakawa, 2001) (Supplemental Figure 2H) and normal numbers of mature B-lymphoid, T-lymphoid, and myeloid lineages in the BM and spleen (Supplemental Figure 2I-J). Hence, I3KO mice demonstrate preserved normal, steady-state adult hematopoiesis, although specific differences in other hematopoiesis conditions need further investigation.

### Igf2bp3 deletion increases the latency of MLL-Af4 leukemia and survival of mice

After confirmation of preserved baseline hematopoiesis in I3KO mice, we next utilized bone marrow transplantation (BMT) assays to query MLL-Af4 mediated leukemogenesis (Supplemental Figure 3A). Retroviral MLL-Af4 transduction was equivalent between WT and I3KO donor BM, based on DNA copy number analysis (Supplemental Figure 3B) and Western blot analysis for MLL-Af4 (Supplemental Figure 3C). Following transplantation of the transduced HSPCs, we found that the loss of *Igf2bp3* significantly increases both the leukemia-free and overall survival of MLL-Af4 mice (Figure 2A-B). The median survival of WT/MLL-Af4 mice was 103 days while I3KO/MLL-Af4 mice had a median survival of greater than 157 days. Complete blood counts of WT/MLL-Af4 mice showed a consistent increase in WBC and absolute myeloid counts over time, which was severely blunted in I3KO/MLL-Af4 mice (Figure 2C; Supplemental Figure 3D). On average, peripheral blood counts crossed the leukemic threshold much earlier in WT/MLL-Af4 mice compared to I3KO/MLL-Af4 mice (70 days versus 112 days) (Figure 2C). Concordantly, peripheral blood smears showed reduced circulating blasts in I3KO/MLL-Af4 mice versus WT/MLL-Af4 mice (Supplemental Figure 3E). Together, these findings indicated that *Igf2bp3* is required for efficient MLL-Af4-mediated leukemogenesis.

**Figure 2:**
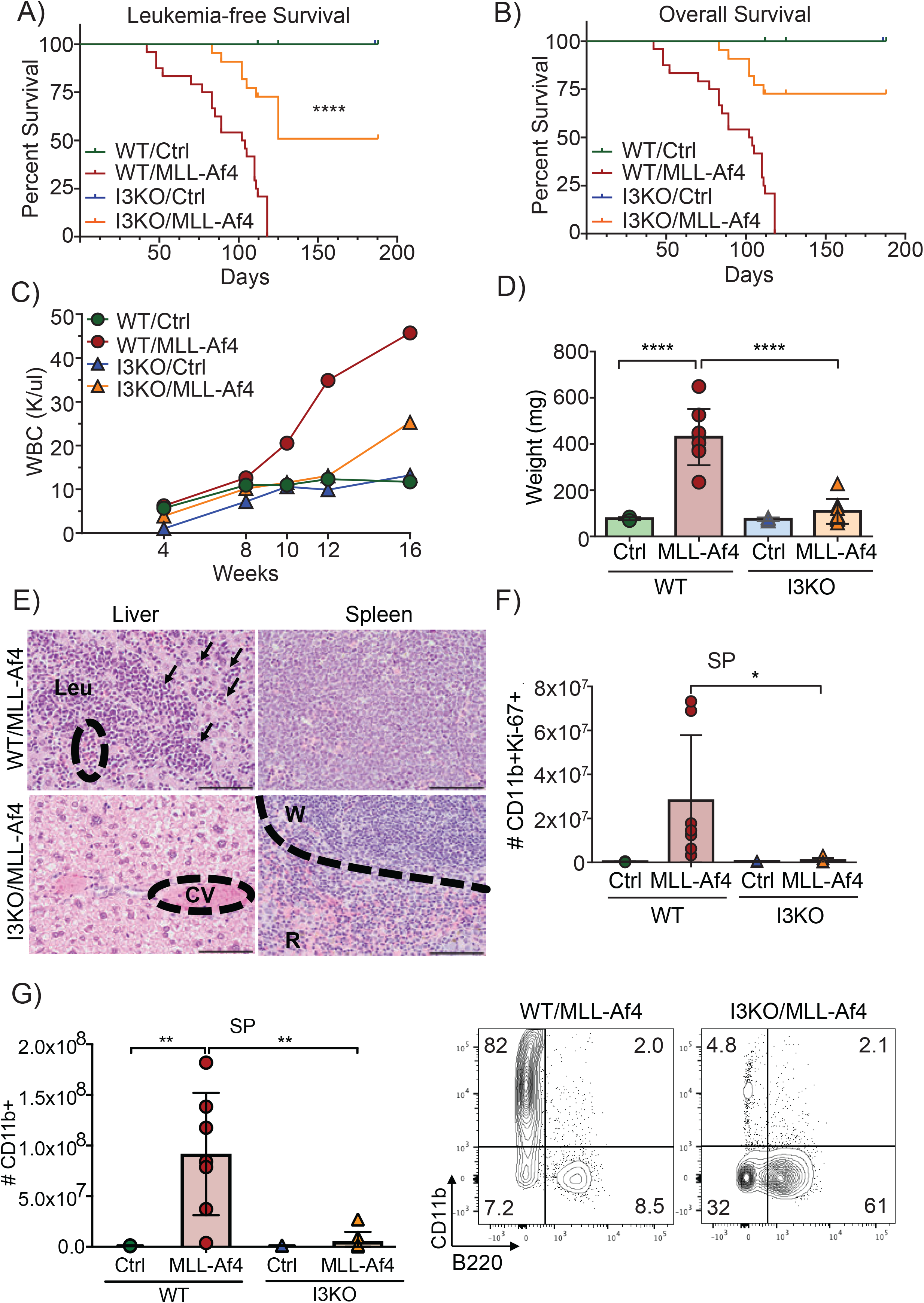
*Igf2bp3* deletion delays leukemogenesis and reduces disease severity. A) Leukemia-free survival of mice transplanted with control (Ctrl) or MLL-Af4 transduced HSPCs from WT or Igf2bp3 KO mice (Kaplan-Meier method with Log-rank test; ****P < 0.0001). B) Overall survival of mice transplanted with Ctrl or MLL-Af4 transduced HSPCs from WT or I3KO mice (n=12 WT/Ctrl, n=24 WT/MLL-Af4, n=7 I3KO/Ctrl, n=22 I3KO/MLL-Af4; Kaplan-Meier method with Log-rank test; ****P < 0.0001). C) Time course of WBC in the PB of mice transplanted with Ctrl or MLL-Af4 transduced HSPCs from WT or I3KO mice (Data represented as means of three experiments; n=4 Ctrl, n=8 MLL-Af4 per experiment). D) Spleen weights of mice transplanted with Ctrl or MLL-Af4 transduced HSPCs from WT or I3KO mice at 14 weeks (n= 4 Ctrl, n=8 MLL-Af4; one-way ANOVA followed by Bonferroni’s multiple comparisons test; ****P < 0.0001). E) H&E staining of liver and spleen of mice transplanted with mice transplanted with MLL-Af4 transduced HSPCs from WT or I3KO mice at 14 weeks. Scale bar:100 microns; CV=Central vein; W=White pulp; R=Red pulp; Leu= Leukemia; arrows showing infiltration. F) Quantitation of CD11b+Ki67+ cells in the spleen at 14 weeks post-transplantation (n= 4 Ctrl, n=8 MLL-Af4; one-way ANOVA followed by Bonferroni’s multiple comparisons test; *P < 0.05). G) (Left) Number of CD11b+ in the SP of recipient mice that received Ctrl or MLL-Af4 transduced HSPCs from WT or I3KO mice at 14 weeks (one-way ANOVA followed by Bonferroni’s multiple comparisons test; **P < 0.01). (Right) Corresponding representative FACS plots showing CD11b+ and B220+ cells in the SP.

### Igf2bp3 modulates disease severity in MLL-Af4-driven leukemia

The MLL-Af4 model utilized here causes a highly penetrant, aggressive form of leukemia in mice. To characterize the role of *Igf2bp3* in disease severity, we performed detailed immunophenotypic and histopathologic analyses in MLL-Af4-transplanted mice in timed experiments. I3KO/MLL-Af4 transplanted mice showed a highly significant, approximately 4-fold reduction in spleen weights at 14 weeks post-transplant compared to WT/MLL-Af4 transplanted mice (Figure 2D). We observed near-total infiltration of the spleen and liver by leukemic cells, obliterating the normal tissue architecture in WT/MLL-Af4 mice, a finding that was much reduced in I3KO/MLL-Af4 mice (Figure 2E). In line with this, I3KO/MLL-Af4 transplanted mice showed a significant reduction in CD11b+ cells (Figure 2G; Supplemental Figure 3G), which were less proliferative (CD11b+Ki-67+), both in the spleen (approximately 30-fold) and in the BM (approximately 2.5-fold) at 14 weeks (Figure 2F; Supplemental Figure 3F). Thus, *Igf2bp3* deletion significantly reduces the tumor burden and attenuates disease severity in MLL-Af4 transplanted mice.

### Igf2bp3 is required for LIC function in vitro

Several studies highlight the importance of LICs in both human and mouse leukemia. In the MLL-Af4 model, these LICs show expression of CD11b and c-Kit (Lin et al., 2017; Somervaille and Cleary, 2006; Somervaille et al., 2009). Given our findings of delayed initiation and decreased disease severity, we characterized these LICs, and found that I3KO/MLL-Af4 transplanted mice showed a significant 10-fold decrease in the numbers of leukemia-initiating cells (CD11b+c-Kit+) in the spleen and a 5-fold decrease in the BM at 14 weeks compared to WT/MLL-Af4 mice (Figure 3A-B). To further characterize the MLL-Af4 LIC and its dependence on IGF2BP3, we turned to endpoint colony formation assays. Utilizing immortalized HSPCs, denoted as Lin-, from WT/MLL-Af4 and I3KO/MLL-Af4 mice, we confirmed of equal transcript expression levels of MLL-Af4 and deletion of IGF2BP3 at the protein level (Figure 3C; Supplemental Figure 5A-B). The deletion of *Igf2bp3* resulted in an approximately 2-fold reduction in total colony formation as well as a significant decrease in CFU-GM progenitors (Figure 3D). To confirm our findings regarding LICs, we utilized an orthogonal method to delete *Igf2bp3* via CRISPR-Cas9. Briefly, Lin-cells from Cas9-GFP BL/6 mice were collected and transduced with the retroviral MSCV-MLL-Af4 vector. After selection, these MLL-Af4 Cas9-GFP Lin-cells were transduced with a retroviral vector with an mCherry fluorescence marker containing either a non-targeting (NT) sgRNA or a sgRNA targeted against *Igf2bp3* (I3sg) (Figure 3E). Importantly, *Igf2bp3* is deleted after transformation with MLL-Af4, a distinction from the previous method. After confirmation of *Igf2bp3* deletion (Figure 3F-G), GFP+mCherry+ MLL-Af4 Lin-cells were utilized for colony-forming assays. We confirmed that CRISPR-Cas9 mediated deletion of *Igf2bp3* led to a significant reduction in total colony numbers and decreases in the various colony morphologies (Figure 3H). The observed differences in overall colony forming capacity between the two systems are most likely a result of the different methodologies being used, but in both systems, IGF2P3 deficiency led to decreased colony formation. Thus, *Igf2bp3* is required for MLL-Af4 LIC function *in vitro*.

**Figure 3:**
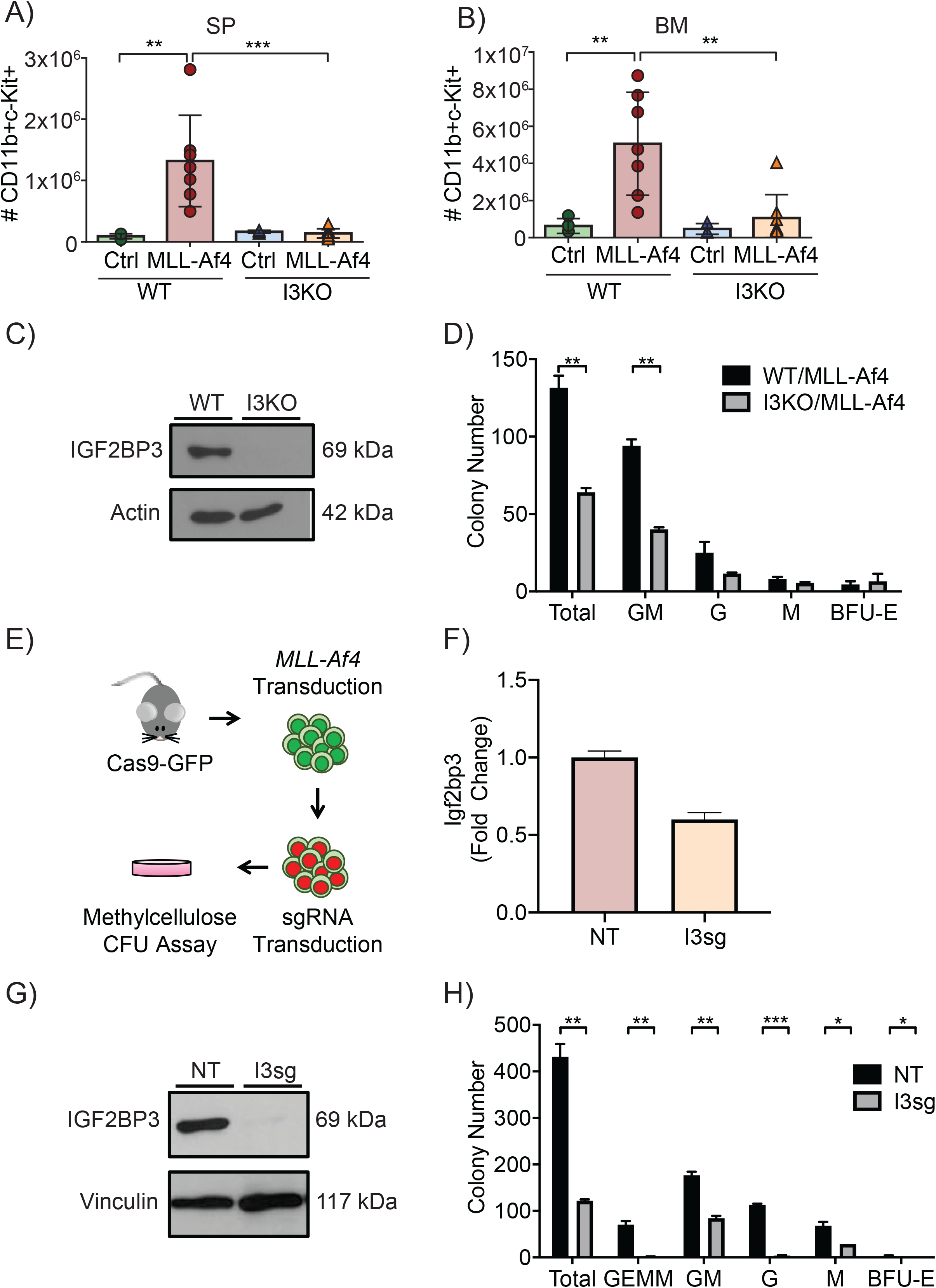
*Igf2bp3* is required for LIC function in endpoint colony formation assays. A) Quantification of CD11b+c-Kit+ cells in the spleen of recipient mice at 14 weeks post-transplantation (n= 4 Ctrl, n=8 MLL-Af4; one-way ANOVA followed by Bonferroni’s multiple comparisons test; **P < 0.01). B) Quantitation of CD11b+c-Kit+ cells in the BM 14 weeks post-transplantation (n= 4 Ctrl, n=8 MLL-Af4; one-way ANOVA followed by Bonferroni’s multiple comparisons test; **P < 0.01, ***P < 0.001). C) Expression of IGF2BP3 of in WT/MLL-Af4 and I3KO/MLL-Af4 immortalized Lin-cells at the protein level. D) Colony formation assay of WT/MLL-Af4 and I3KO/MLL-Af4 immortalized Lin-cells (t-test; **P < 0.01). E) Schematic of collection of Cas9-GFP MLL-Af4 Lin-cells and CRISPR-Cas9 mediated deletion of *Igf2bp3*. F) Expression of Igf2bp3 in Cas9-GFP MLL-Af4 Lin-cells in non-targeting (NT) and Igf2bp3 deleted (I3sg) cells by RT-qPCR. G) Expression of IGF2BP3 in NT and I3sg Cas9-GFP MLL-Af4 Lin-cells at the protein level. H) Colony formation assay of NT and I3sg deleted Cas9-GFP MLL-Af4 Lin-cells (t-test; *P < 0.05, **P < 0.01, ***P <0.001).

### Igf2bp3 is necessary for the function of MLL-Af4 leukemia-initiating cells in vivo

Since *Igf2bp3* deletion causes a reduction in LICs and is required for the function of these LICs, we next wanted to determine if *Igf2bp3* specifically affects the capability of these LICs to initiate MLL-Af4 leukemia *in vivo*. First, to investigate baseline hematopoietic stem cell function in I3KO mice, we completed a competitive repopulation bone marrow transplantation experiment by transplanting lethally irradiated CD45.1 recipient mice with 50% of either WT or I3KO CD45.2 donor BM and 50% CD45.1 donor BM. We found no defect in engraftment over time in transplant recipients of I3KO BM (Supplemental Figure 4A). Moreover, we determined no differences in multilineage hematopoietic reconstitution ability of I3KO donor cells, as immature lineages in the BM and mature B-hematopoietic cells in the periphery were intact (Supplemental Figure 4B-H, respectively). Given that there were no baseline differences in reconstitution by normal HSPCs, we now sought to determine if *Igf2bp3* impacted the number of effective LICs in secondary transplant assays. We isolated BM cells from WT/MLL-Af4 and I3KO/MLL-Af4 mice with equivalent disease burdens and transplanted equal numbers (10^6^, 10^5^, and 10^4^) of leukemic BM cells into immunocompetent CD45.1 mice. At 4 weeks post-transplantation, mice that received 10^6^ cells from I3KO/MLL-Af4 mice had significantly reduced donor CD45.2+ cell engraftment (Figure 4A). With 10^5^ and 10^4^ transplanted cells, we no longer observed measurable leukemic burden in recipient mice (Figure 4A). This suggests that the active cell frequency of LICs in I3KO/MLL-Af4 mice is lost between 10^6^ and 10^5^ cells (Figure 4A)(Brien et al., 2010). Moreover, WBC and splenic weights were significantly decreased in I3KO/MLL-Af4 leukemia transplanted mice (Figure 4B-D). Histologically, leukemic infiltration was absent in the spleen and liver of 10^5^ transplanted I3KO/MLL-Af4 mice (Figure 4E). Thus, *Igf2bp3* deletion greatly attenuated transplantability, in which only 17% of I3KO/MLL-Af4 recipients developed leukemia while 67% of WT/MLL-Af4 recipients developed leukemia at 4 weeks with 10^6^ transplanted cells (Figure 4B). These data show that the deletion of *Igf2bp3* results in the significant reduction of LICs and reconstitution of MLL-Af4 transplanted mice, suggesting that *Igf2bp3* is necessary for the self-renewal capability of LICs *in vivo*.

**Figure 4:**
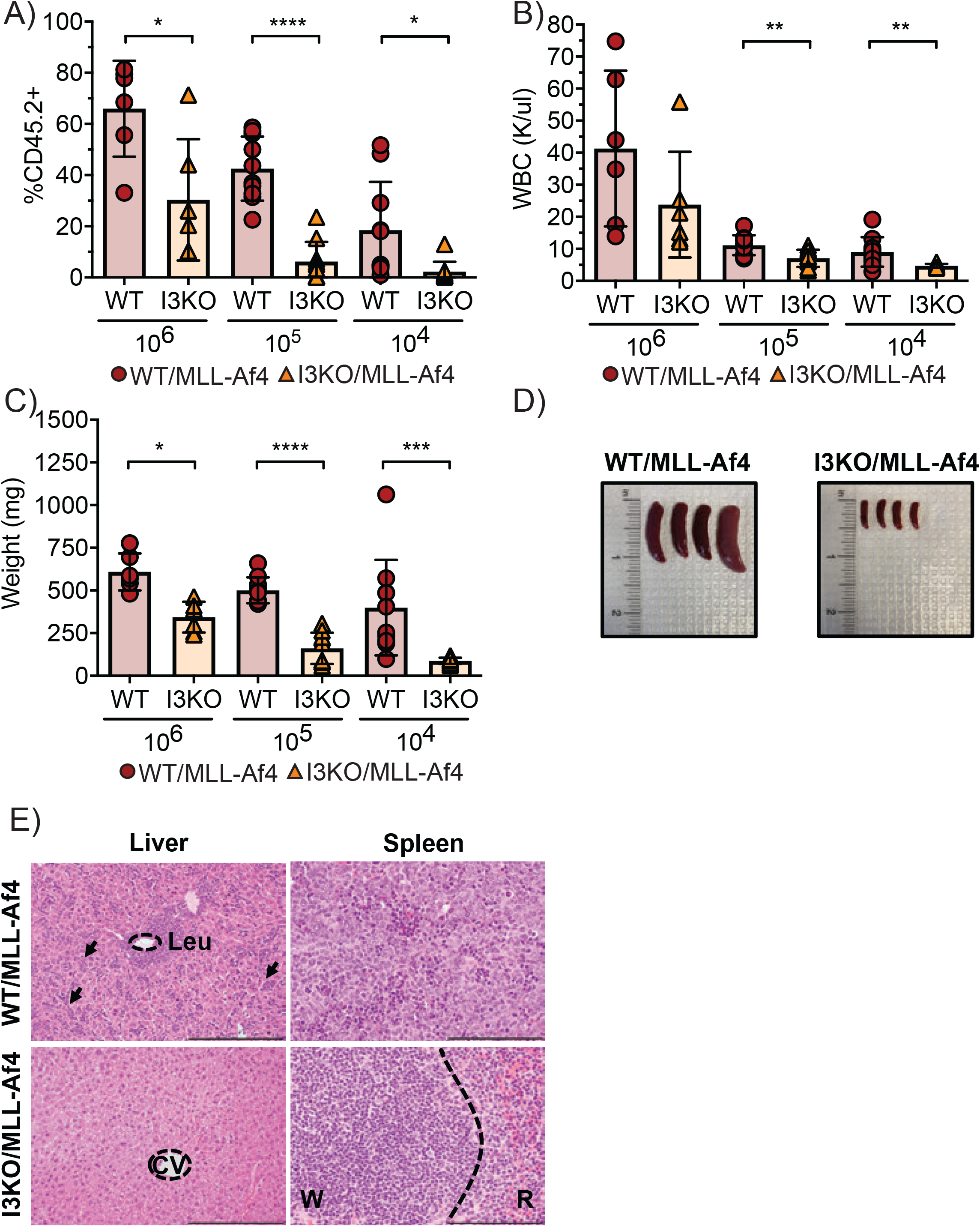
*Igf2bp3* deletion is necessary for MLL-Af4 leukemia-initiating cells to reconstitute mice *in vivo*. A) Percentage of CD45.2+ in the peripheral blood of secondary transplanted mice from leukemic WT/MLL-Af4 or I3KO/MLL-Af4 donor mice at 10^6^, 10^5^, and 10^4^ BM cells at 4 weeks post-transplantation (n= 6 10^6^, n=10 10^5^, n=10 10^4^; t-test; *P < 0.05, ***P < 0.001, ****P < 0.0001). B) WBC from PB of secondary transplanted mice from WT/MLL-Af4 or I3KO/MLL-Af4 BM 3-4 weeks post-transplant (n= 6 10^6^, n=10 10^5^, n=10 10^4^; t-test; **P < 0.01). C) Splenic weights of secondary transplanted mice at 4-5 weeks (n= 6 10^6^, n=10 10^5^, n=10 10^4^; t-test; *P < 0.05, ***P < 0.001, ****P < 0.0001). D) Images of splenic tumors in secondary mice transplanted with 10,000 BM cells from WT/MLL-Af4 mice (left) or I3KO/MLL-Af4 mice (right) at 5 weeks. E) H&E staining of liver and spleen of secondary transplant recipients that received 10^5^ cells at 4 weeks. Scale bar: liver, 200 microns; spleen, 100 microns; CV=Central vein; W=White pulp; R=Red pulp; Leu= Leukemia; arrows showing infiltration.

### IGF2BP3 supports oncogenic gene expression networks in LIC-enriched and bulk leukemia cells

To identify differentially expressed transcripts related to the I3KO phenotype, we sequenced RNA from WT/MLL-Af4 and I3KO/MLL-Af4 Lin- and CD11b+ bulk leukemia cells (Figure 5A-B, respectively). First, we confirmed expression of MLL and *Igf2bp3* in these samples by RT-qPCR and WB (Figure 3C; Supplemental Figure 5A-E). Differential expression analysis by DEseq2 revealed hundreds of differentially expressed transcripts (Figure 5A-B; Tables S2 and S3) (Love et al., 2014). We observed 208 upregulated and 418 downregulated transcripts in the CD11b+ cells, and 189 upregulated and 172 downregulated transcripts in the Lin-cells. To identify over-represented pathways and gene ontology terms within IGF2BP3 differentially regulated transcripts, we used the Metascape analysis tool (Zhou et al., 2019) and observed a significant enrichment in transcripts associated with the KEGG term related to transcriptional misregulation in cancer in both the Lin- and CD11b+ bulk leukemia dataset (Figure 5C-D). Interestingly, there were also distinct oncogenic networks that were regulated in the two datasets, with regulation of discrete signaling pathways noted in the Lin-cells (PI3K/AKT) and in the CD11b+ cells (GTPase, MAPK pathway) (Figure 5C-D). This was also confirmed by an independent analysis of differentially expressed data using GSEA, where we noted that the Hallmark KRAS pathway was significantly enriched in the CD11b+ cells (Supplemental Figure 5G) and GO oxidative phosphorylation in the Lin-cells (Supplemental Figure 5H). We used RT-qPCR to validate the RNA-seq data from both Lin- and CD11b+ cells. In Lin-cells, we focused on differentially regulated genes with a known leukemogenic function including *Csf2rb, Notch1, Cd69*, and the *Hoxa* cluster of transcripts, including *Hoxa9, Hoxa10, Hoxa7*. We observed a significant decrease in the steady state mRNA levels for each of these transcripts in the I3KO/MLL-Af4 Lin-cells, confirming our RNA sequencing results (Figure 5E). In CD11b+ cells, we selected transcripts known to play a role in Ras signaling, *Ccnd1, Maf, Mafb Itga6, Klf4, and Akt3* (Brundage et al., 2014; Eychène et al., 2008; Lewis et al., 2019; Riverso et al., 2017; Takata et al., 2005; Wagle et al., 2018; Wu et al., 2019). As expected, these transcripts were decreased in I3KO/MLL-Af4 CD11b+ cells by RT-qPCR, confirming the high throughput RNA sequencing findings (Figure 5F). Furthermore, we determined that there was a significant decrease in Ras GTPase activity in the I3KO/MLL-Af4 CD11b+ cells compared to WT/MLL-Af4 CD11b+ bulk leukemia cells by ELISA assay (Figure 5G). Together, this data demonstrates that IGF2BP3 plays a major role in amplifying the expression of many cancer-related genes in Lin- and CD11b+ cells.

**Figure 5:**
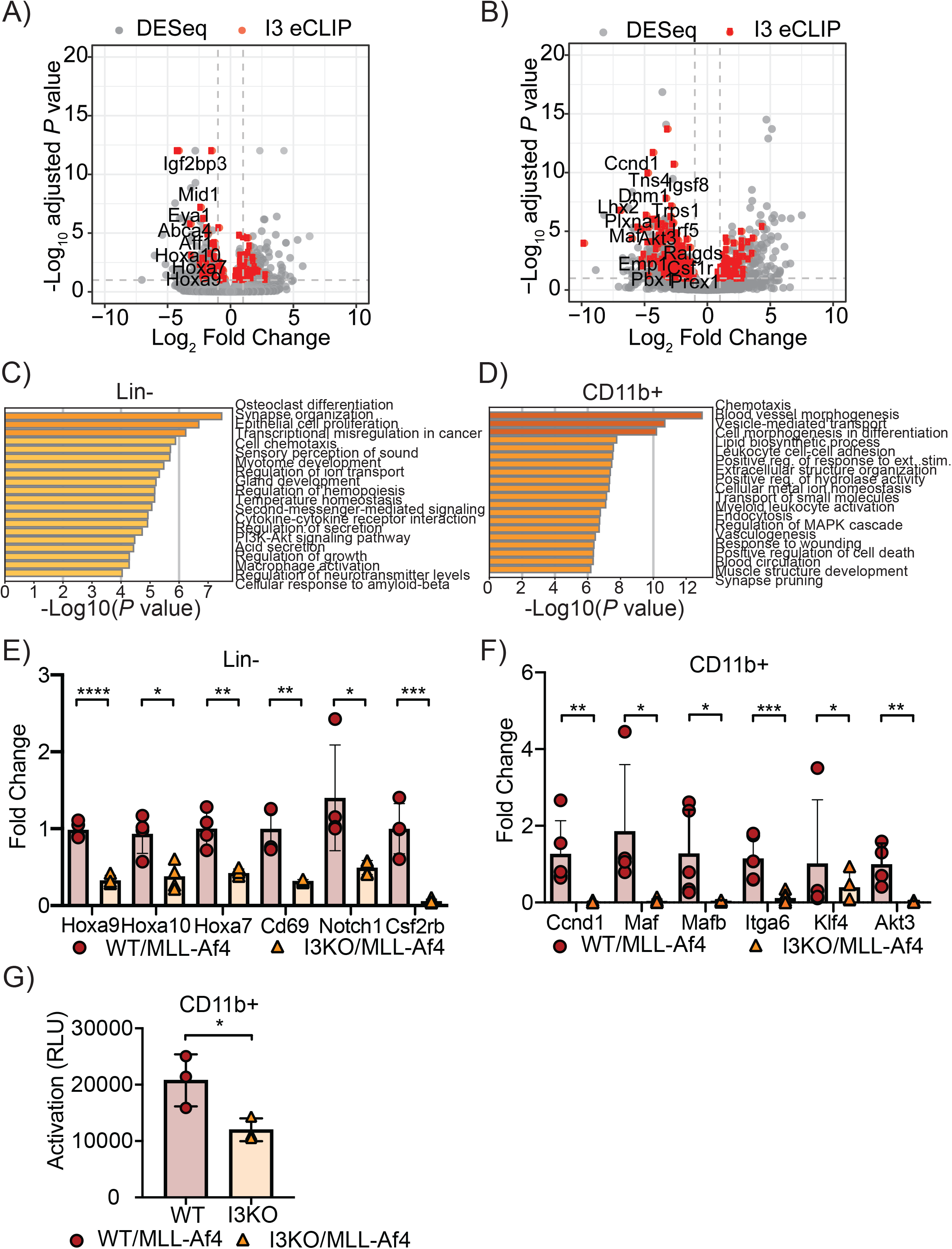
IGF2BP3 enhances MLL-Af4 mediated leukemogenesis through targeting transcripts within leukemogenic and Ras signaling pathways. A) Volcano plot of differentially expressed genes determined using DESeq analysis on RNA-seq samples from WT/MLL-Af4 or I3KO/MLL-Af4 Lin-cells. Dotted lines represent 1.0-fold–change in expression (vertical lines) and adjusted *P* < 0.1 cutoff (horizontal line). IGF2BP3 eCLIP-seq targets are highlighted in red. B) Volcano plot of differentially expressed transcripts determined using DESeq analysis on RNA-seq samples from WT/MLL-Af4 or I3KO/MLL-Af4 CD11b+ cells. Dotted lines represent 1.0-fold–change in expression (vertical lines) and adjusted *P* < 0.1 cutoff (horizontal line). IGF2BP3 eCLIP-seq targets are highlighted in red. C) GO Biological Processes and KEGG Pathway enrichment determined utilizing the Metascape enrichment analysis webtool on MLL-Af4 Lin-IGF2BP3 DESeq dataset with an adjusted P < 0.05 cutoff. D) GO Biological Processes and KEGG Pathway enrichment determined utilizing the Metascape enrichment analysis webtool on MLL-Af4 CD11b+ IGF2BP3 DESeq dataset with an adjusted P < 0.05 cutoff. Bar graphs are ranked by P value and overlap of terms within gene list. E) Expression of leukemogenic target genes in WT/MLL-Af4 and I3KO/MLL-Af4 Lin-cells by RT-qPCR (n= 4; t-test; *P < 0.05, **P < 0.01, ****P < 0.0001). F) Expression of Ras signaling pathway genes in WT/MLL-Af4 and I3KO/MLL-Af4 CD11b+ cells by RT-qPCR (n=4; t-test; *P < 0.05, **P < 0.01, ***P < 0.001). G) Ras GTPase activity by ELISA in WT/MLL-Af4 and I3KO/MLL-Af4 CD11b+ cells (n=3; t-test; *P < 0.05).

### eCLIP analysis reveals a putative role for IGF2BP3 in pre-mRNA splicing

To determine how IGF2BP3 modulates gene expression in MLL-Af4 leukemia, we performed eCLIP to identify IGF2BP3 bound transcripts in both Lin- and CD11b+ cells (Figure 5A-B; eCLIP target transcripts denoted in red). Sequencing libraries were prepared from a minimum of two biological replicates with two technical replicates (four immunoprecipitations per cell line) as well as size matched input (smInput) samples from each condition. After filtering out reads that overlap the smInput, reproducible peaks were identified by using CLIPper (Tables S2-3; FS1 column) (Lovci et al., 2013). A significant fraction of the differentially expressed mRNAs are bound by IGF2BP3 (P< 2.2×10^−^16; Supplemental Figure 6A). Motif analysis revealed an enrichment of CA-rich elements as expected (Supplemental Figure 6B) (Schneider et al., 2019). Although the majority of peaks were present within introns, we observed cell type-specific differences in the locations of IGF2BP3 binding sites within exons. In CD11b+ cells, a greater proportion of exonic peaks were found in 3’UTRs whereas a greater proportion of peaks mapped to internal exons in Lin-cells (Figure 6A). The eCLIP data revealed numerous peaks within precursor mRNA (pre-mRNA) in both Lin- and CD11b+ cells, suggesting a potential role in splicing regulation. To characterize this observation, we utilized MISO (Mixture of Isoforms) analysis to identify differentially spliced transcripts in WT/MLL-Af4 and I3KO/MLL-Af4 cells (Katz et al., 2010). Across both cell lines, we identified hundreds of transcripts with IGF2BP3-dependent changes in alternative splicing, including 97 differential splicing events in Lin- and 261 splicing events in CD11b+ cells (Bayes factor ≥ 10, delta PSI ≥ 0.1, and minimum 20 reads supporting the event) (Supplemental Figure 6C). After merging all replicate eCLIP data for each cell type, we determined the position of eCLIP peaks relative to splice sites for splicing events identified by MISO (Figure 6B). Most event types, including skipped exons (SE), alternative first exons (AFE), alternative last exons (ALE), alternative 3’ splice sites (A3SS), and alternative 5’ splice sites (A5SS) exhibited both increases and decreases in percent spliced in (PSI), however, intron retention (RI) events showed a consistent reduction in splicing in the I3KO/MLL-Af4 cells (Figure 6C). A significant fraction of alternatively spliced transcripts contained IGF2BP3 binding sites in proximity of the splicing event (P<2.2×10^−16^, Supplementary Figure 6D). Across all event types we found that the density of IGF2BP3 binding sites was strongest near the 3’splice site (3’ss), with additional signal near the 5’ splice site (5’ss). This pattern was observed for each distinct splicing event class that MISO identified, with retained introns exhibiting the strongest bias towards the 3’ss (Supplemental Figure 6E). This positional bias in the data was noted for some differentially expressed genes, such as *Hoxa9, Hoxa7*, and *Cd69* (Figure 6D). *Hoxa9* is known to be alternatively spliced, with two well-characterized transcripts, full-length *Hoxa9* and truncated (homeobox-less) *Hoxa9T* (He et al., 2012; Stadler et al., 2014). We designed and validated RT-qPCR primers to measure the two RNA isoforms, finding that I3KO/MLL-Af4 cells showed an alteration in the ratio of the two isoforms (Figure 6F). This was in addition to our previous finding that the total transcript (using primers that are isoform-agnostic) was also downregulated in I3KO/MLL-Af4 cells (Figure 5E). Hence, the net effect of IGF2BP3 may be multi-pronged—there is a strong impact on steady state mRNA levels and potentially an impact on splicing. Taken together, these data demonstrate that IGF2BP3 functions in regulation of alternative pre-mRNA splicing in bulk leukemia cells and progenitor cells.

**Figure 6:**
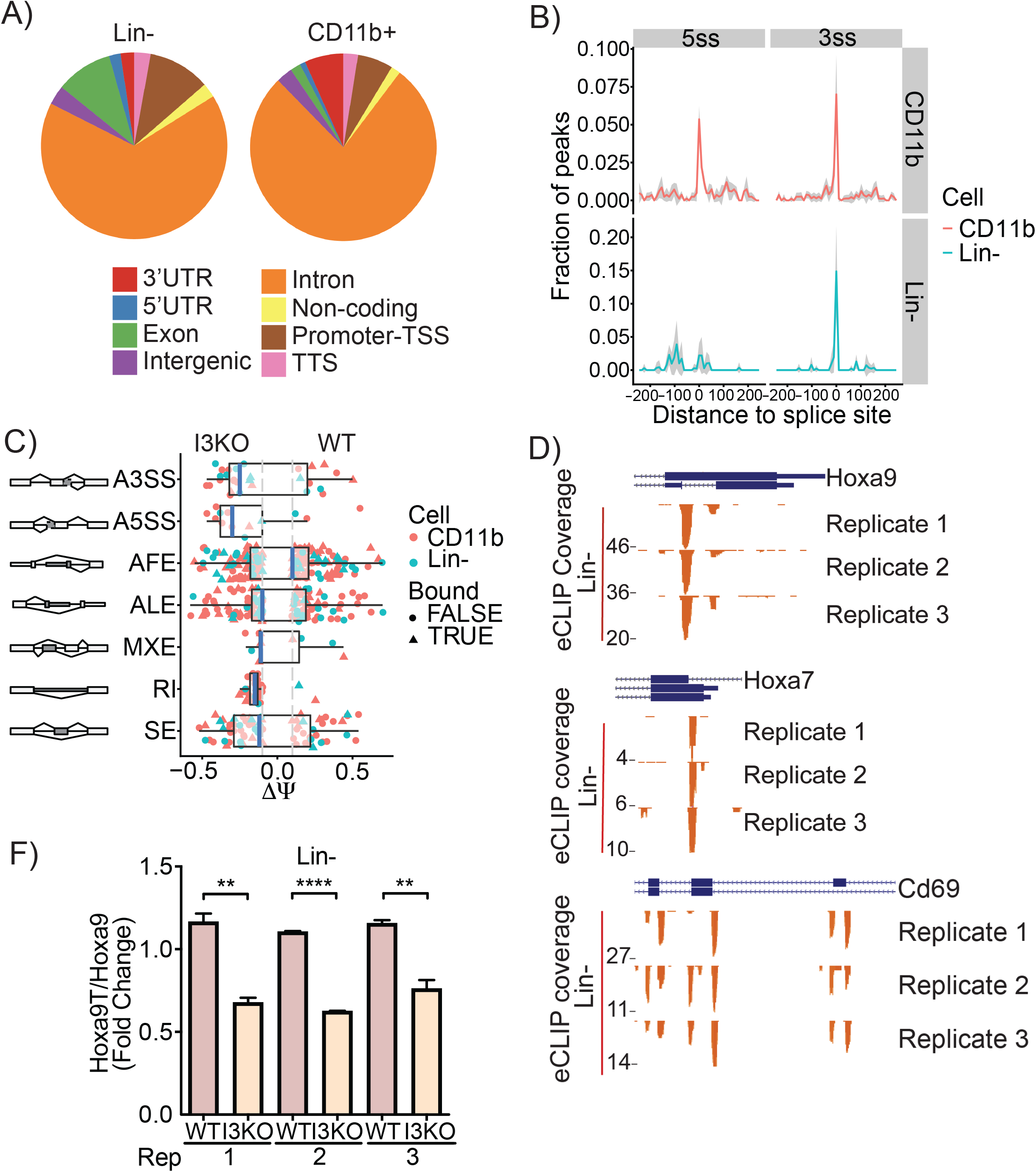
eCLIP analysis reveals IGF2BP3 function in regulating alternative pre-mRNA splicing. A) Genomic locations of IGF2BP3 eCLIP peaks in WT/MLL-Af4 Lin-cells and CD11b+ cells. B) Histogram showing normalized IGF2BP3 eCLIP peak counts and distance from IGF2BP3 eCLIP peak of 5’ (5ss) and 3’ (3ss) splice sites in WT/MLL-Af4 CD11b+ (top) cells and Lin-cells (bottom). C) Distribution of types of alternative splicing patterns for WT/MLL-Af4 or I3KO/MLL-Af4 Lin- and CD11b+ cells using MISO analysis. Delta psi values plotted indicate difference in isoforms.(A3SS, Alternative 3’ splice sites; A5SS, Alternative 5’ splice sites; AFE, Alternative first exons; ALE, Alternative last exons; MXE, Mutually exclusive exons; RI, Retained introns; SE, Skipped exons; Bound, IGF2BP3 eCLIP target). D) UCSC Genome Browser snapshots of the Hoxa9, Hoxa7, and Cd69 loci. Each panel shows the exon-intron structure of the gene and unique read coverage from 3 eCLIP biological replicates from WT/MLL-Af4 Lin-cells. The maximum number of reads at each position is indicated to the left of each histogram. F) Ratio of Hoxa9T/Hoxa9 isoforms in WT/MLL-Af4 and I3KO/MLL-Af4 Lin-cells by RT-qPCR (t-test; **P < 0.01, ****P < 0.0001).

## DISCUSSION

Here, we have generated an *Igf2bp3*–deficient murine model and queried MLL-Af4 mediated leukemogenesis. We demonstrated that *Igf2bp3* is required for the efficient initiation of leukemia and that this regulates the number and frequency of MLL-Af4 LICs. *Igf2bp3* regulates the expression of numerous critical transcripts in the *Hoxa* locus and the Ras signaling pathway, leading to dysregulated gene expression and enhanced downstream signaling, thereby promoting leukemogenesis.

MLL-AF4 driven leukemogenesis is associated with massive transcriptional dysregulation, mediated by the fusion of a histone methyltransferase with a factor involved in the super elongation complex (Smith et al., 2011). We confirm here that *Igf2bp3* is a direct transcriptional target of the MLL-AF4 fusion protein. Interestingly, IGF2BP3 itself seems to positively regulate MLL-AF4 transcriptional targets, based on our analysis provided here. Together, these data suggest that IGF2BP3 forms a novel post-transcriptional feed-forward loop that stabilizes and/or enhances the expression of MLL-Af4 transcriptional targets. Because of this unique relationship, and its relatively restricted pattern of expression in MLL-translocated leukemia, it is not clear if IGF2BP3 may play a role in other leukemia subtypes. However, it is worth noting that IGF2BP3 overexpression is noted in a wide range of cancer types and, hence, additional work is required to establish its role in other types of hematologic and non-hematologic malignancy.

In our previous study, we determined that IGF2BP3 is required for B-ALL cell survival, and that overexpression of IGF2BP3 in the bone marrow of mice leads to a pathologic expansion of hematopoietic stem and progenitor cells (Palanichamy et al., 2016). Here, using the MLL-Af4 leukemia model, we found that the deletion of *Igf2bp3* caused a striking delay in leukemia development and significantly increased the survival of MLL-Af4 mice. Furthermore, *Igf2bp3* deficiency greatly attenuated the aggressiveness of the disease. This was demonstrated by significant decreases in WBC counts, spleen weights, and infiltrating leukemic cells visualized in histopathological analysis of hematopoietic tissues. Although MLL-Af4 drives an acute myeloid leukemia in mice (Lin et al., 2016), it is important to note that *MLL*r leukemias often show lineage infidelity and plasticity, leading to difficulties in applying targeted therapy (Rayes et al., 2016). While our prior work focused on IGF2BP3 in B-lineage *MLL*r leukemia, the current work suggests its broader applicability to all *MLL*r leukemia. Hence, IGF2BP3 may be a constant factor to target across the phenotypic range of *MLL*r leukemia and may be less subject to change in response to targeted therapy.

We also determined that *Igf2bp3* regulates the numbers and function of leukemia-initiating cells (LICs). Importantly, the effect of *Igf2bp3* deletion was restricted to LICs and did not significantly impact normal HSC function. Deletion of *Igf2bp3* led to a LIC disadvantage *in vivo* and *in vitro*, using both the I3KO mouse and a novel, orthogonal system utilizing CRISPR/Cas9-mediated deletion of *Igf2bp3*. LICs have been defined as cells that can self-renew and have the capability to produce downstream bulk leukemia cells (Magee et al., 2012). The persistence of these LICs is thought to contribute to relapse after treatment in several different leukemia subtypes (Bao et al., 2006; Chu et al., 2011; Diehn et al., 2009; Merlos-Suárez et al., 2011). In *MLL*r leukemia, LICs have been shown to have a high frequency in tumors and co-expression of mature lineage-restricted cell markers, with some excellent work in mouse models (Krivtsov et al., 2006; Somervaille and Cleary, 2006; Somervaille et al., 2009). However, the details of human LICs, particularly in MLL-AF4 leukemia, are less well known (Agraz-Doblas et al., 2019; Bardini et al., 2015; Barrett et al., 2016; Metzler et al., 2006). The role of IGF2BP3 in such cells will be of great interest and is a future direction for our work.

Previously, we discovered that IGF2BP3 interacts primarily with the 3’UTR of its target transcripts via iCLIP-seq (Palanichamy et al., 2016). In this study, we determined that IGF2BP3 targets many transcripts within intronic regions and near splice sites in addition to the 3’UTR, suggesting additional roles in post-transcriptional gene regulation. This difference may be due to the use of the eCLIP technique or the focused application on primary cells as opposed to cell lines. It is not entirely surprising, however, since RBPs are known to regulate gene expression at several steps at the post-transcriptional level through mRNA operons (Keene, 2007; Keene and Lager, 2005). Furthermore, a recent study has shown that IGF2BP3 may regulate alternative splicing in the PKM gene in lung cancer cells (Xueqing et al., 2020). In line with this study, we found dynamic alternative splicing events that reflected various categories of alternative splicing phenomena, including retained Introns, alternative 5’ and 3’ splice sites, and skipped exons. In this light, it is interesting to note that intron retention has recently been reported to be a mechanism of transcriptome diversification in cancer and, specifically, in leukemia (Dvinge and Bradley, 2015; Wang et al., 2019). With this unexpected, novel discovery and our prior work detailing interactions with the 3’UTR, it is very likely that IGF2BP3 regulates specific mRNA operons through multiple post-transcriptional mechanisms in MLL-Af4 driven leukemia.

As an RBP, the function of IGF2BP3 is intimately connected to the underlying transcriptional program—IGF2BP3 can only act on transcripts that are specifically induced in the cell type where it is expressed. Hence, the finding of unique sets of genes that are bound and regulated by IGF2BP3 in Lin- and CD11b+ cells is not entirely unexpected, given that transcription changes as the leukemia initiating cells differentiate into the bulk leukemic cells. This is similar to what has been observed for miRNAs, which post-transcriptionally regulate distinct gene expression programs in distinct cell types (Lechman et al., 2016). The significant enrichment of IGF2BP3-bound mRNAs in the sets of differentially regulated and differentially spliced transcripts confirms a direct regulatory effect. However, further work is required to confirm functional relationships between the specific transcripts that are regulated and the phenotypic effects driven by IGF2BP3.

Notably, these differentially regulated transcripts showed significant enrichment for the KEGG transcriptional misregulation in cancer term as well as the GO oxidative phosphorylation term. Notable IGF2BP3 targets included critical transcripts in the *Hoxa* cluster such as *Hoxa9, Hoxa10, and Hoxa7*. HOXA9 is induced by MLL-AF4, plays a role in normal hematopoiesis, and is required for the survival of MLL-rearranged leukemia (Ernst et al., 2004b; Faber et al., 2009; Imamura et al., 2002; Lawrence et al., 2005; Pineault et al., 2002; Rozovskaia et al., 2001). Furthermore, Hoxa9 is an alternatively spliced gene, with co-expression of a homeodomain-less splice variant, *Hoxa9T*, together with *Hoxa9*, shown to be necessary for full leukemogenic transformation (He et al., 2012; Stadler et al., 2014). Hence, *Igf2bp3* may act through upregulation of *Hoxa9* and *Hoxa9T* through multiple post-transcriptional mechanisms to promote MLL-Af4 driven leukemogenesis and impact the function of MLL-Af4 LICs. In addition, HOXA9 may play a role in the regulation of oxidative phosphorylation (Lynch et al., 2019), and it is tempting to speculate that the observed *Igf2bp3*-dependent impact on LICs is a consequence of dysregulated oxidative phosphorylation, a key pathway that regulates LICs. Importantly, because *Igf2bp3* was not required for steady-state hematopoiesis, in contrast to HOXA9, it may represent a more attractive target.

Work from our lab and others have demonstrated that IGF2BP3 targets a wide array of oncogenic transcripts and pathways, including CDK6 and MYC (Palanichamy et al., 2016). Here, we found that IGF2BP3 targets and modulates the expression of many transcripts within the Ras signaling pathway and its downstream effector pathways. RAS proteins control numerous cellular processes such as proliferation and survival, and are amongst the most commonly mutated genes in cancer (Downward, 2003; Schubbert et al., 2007). Interestingly, while *MLL*r leukemia has a paucity of additional mutations, the mutations that are present are in found mainly in the RAS signaling pathway (Agraz-Doblas et al., 2019; Andersson et al., 2015; Chandra et al., 2010; Emerenciano et al., 2015; Hyrenius-Wittsten et al., 2018; Kerstjens et al., 2017; Lavallée et al., 2015; Trentin et al., 2016). Moreover, several studies have shown selective activity against MLL-r leukemia cell lines and primary samples *in vitro* by MEK inhibitors, suggesting an important role for signaling downstream of RAS mutations in leukemia cell survival and proliferation (Kampen et al., 2014; Kerstjens et al., 2017; Lavallée et al., 2015).

Here, we determined that *Igf2bp3* is required for the efficient initiation of MLL-Af4 driven leukemia as well as for the development of and self-renewal capability of MLL-Af4 LICs. Mechanistically, IGF2BP3 binds to hundreds of transcripts and modulates their expression *in vivo* and *in vitro* through multiple, complex post-transcriptional mechanisms. We describe a novel positional bias for IGF2BP3 binding in leukemic cells isolated from an *in vivo* model, a notable advance. In summary, our study demonstrated that IGF2BP3 is an amplifier of *MLL*r leukemogenesis by targeting *Hoxa* transcripts essential for leukemia-initiating cell function and targeting Ras signaling pathway transcripts, thereby controlling multiple critical downstream effector pathways required for disease initiation and severity. Our findings highlight IGF2BP3 as a necessary regulator of MLLr leukemia and a potential therapeutic target for this disease.

## METHODS

### ChIP-PCR

RS4;11 and SEM cells were used for ChIP assays as previously described (Janardhan et al., 2017). Primer sequences for the IGF2BP3 promoter region were provided by James Mulloy (University of Cincinnati College of Medicine)(Lin et al., 2016).

### Western Blotting and RT-Qpcr

Protein and mRNA extracts were prepared, and Western Blot/RT-qPCR performed as previously described (Fernando et al., 2017). Primers for qPCR and antibodies used for Western blotting are listed in Table S4.

### Plasmids

The MSCV-MLL-flag-*Af4* plasmid was kindly provided by Michael Thirman (University of Chicago, Department of Medicine) through MTA (Lin et al., 2016). The non-targeting or *Igf2bp3* sgRNA was cloned into an in-house MSCV-hU6-sgRNA-EFS-mCherry vector.

### Retroviral transduction and bone marrow transplantation

Retroviral transduction and bone marrow transplantation (BMT) were completed as previously described (Fernando et al., 2017; O’Connell et al., 2010; Rao et al., 2010). 5-FU enriched BM and Lin-cells were spin-infected four times with MSCV-MLL-flag-*Af4* virus at 30°C for 45 minutes in the presence of polybrene. Cells were selected with 400 μg/ml of G418 for 7 days. For sgRNA-mediated knockout, MLL-Af4 overexpressing Cas9-GFP cells were retrovirally infected with MSCV-hU6-NT/I3sgRNA-EFS-mCherry. The selected cells were then cultured for colony formation assays or injected into lethally irradiated mice.

### Mice

C57BL/6J and B6J.129(Cg)-Gt(ROSA)26Sor^tm1.1(CAG-cas9*,-EGFP)Fezh^/J (Cas9-GFP) mice were obtained from Jackson Laboratory. For *Igf2bp3* KO mouse generation, the UCI Transgenic Mouse Facility utilized CRISPR/Cas9 to insert loxP sites flanking exon 2 of *Igf2bp3* to generate *Igf2bp3*^f/f^ mice. We originally attempted to generate conditional KO mice by breeding the *Igf2bp3*^f/f^ mice with Vav1-Cre mice. Consistent with prior reports, we found that this strategy led to “leaky” Cre expression, resulting in germline deletion (Croker et al., 2004; de Boer et al., 2003; Georgiades et al., 2002; Heffner et al., 2012; Joseph et al., 2013). To isolate the floxed and deletion (del) alleles, we back-crossed the mice onto C57BL/6 mice, successfully confirming germline, Mendelian transmission of the del and floxed alleles in two successive generations (Table S1). Mice heterozygous for the del allele were mated together, with the production of a homozygous deletion of *Igf2bp3*, resulting in the *Igf2bp3*^del/del^ mice (I3KO) used in this study.

### Cell culture

RS4;11, SEM, 70Z/3 and HEK 293T cell lines were cultured as previously described (Fernando et al., 2017). Lin-cells were cultured in IMDM with 15% fetal bovine serum supplemented with SCF, IL-6, FLT3, and TPO. CD11b+ cells were isolated from splenic tumors for positive selection by CD11b antibody and MACS (Miltenyi).

### Flow cytometry

Blood, BM, thymus, and spleen were collected from the mice under sterile conditions at the indicated time points and staining performed as previously described (Contreras et al., 2015; Fernando et al., 2017). The list of antibodies used is provided in Table S4. Flow cytometry was performed on a BD FACS LSRII. Analysis was performed using FlowJo software.

### Histopathology

Fixation and sectioning has been described previously (O’Connell et al., 2010). Analysis was performed by a board certified hematopathologist (D.S. Rao).

### Competitive repopulation assay and secondary leukemia transplantation

Competitive repopulation experiments were completed as previously described (Palanichamy et al., 2016). For leukemia transplantation, BM was collected from WT/MLL-Af4 or I3KO/MLL-Af4 mice that succumbed to leukemia at 10-14 weeks post-transplantation and injected into 8-week-old immunocompetent CD45.1+ female mice.

### eCLIP

IGF2BP3 crosslinking-immunoprecipitation studies were carried out from a minimum of two biological replicates with two technical replicates (four immunoprecipitations per cell type) and size matched input (smInput) samples in each cell type using Eclipse BioInnovations eCLIP kit. Briefly, 5×10^5^ cells were crosslinked with 245nm UV radiation at 400mJoules/cm^2^. Crosslinked cell lysates were treated with RNAse I to fragment RNA and immunoprecipitated with anti-IGF2BP3 antibody (MBL RN009P) coupled to magnetic Protein G beads. Paired-end RNA sequencing was performed on the Illumina HiSeq4000 system at the UCSF Genomics Core Facility. Peaks were called using CLIPper (Lovci et al., 2013). Peaks were filtered based on appearance in the smInput (FS1). Annotation of the genomic location of the peaks and motif enrichment analysis were performed using HOMER (Heinz et al., 2010) annotatePeaks.pl and findMotifs.pl, respectively. Background for the peaks within differentially expressed genes was simulated using bedtools (Heinz et al., 2010; Quinlan and Hall, 2010) and shuffled 1000 times.

### RNA seq

Single-end, strand-specific RNA sequencing was performed on the Illumina HiSeq3000 system for the Lin- and CD11b+ samples, resulting in 15-20 million reads per sample, at the UCLA Technology Center for Genomics & Bioinformatics. Our analysis pipeline has been previously described (Palanichamy et al., 2016). Enrichment analysis for KEGG pathways and Gene Ontology (GO) biological processes terms was completed with the Metascape analysis tool (http://metascape.org)(Zhou et al., 2019). Gene Set Enrichment Analysis (GSEA) was completed using the GSEAPreranked software on both the Lin- and CD11b+ DESeq datasets after calculation of π-value (Mootha et al., 2003; Subramanian et al., 2005; Xiao et al., 2014) to compare to the Hallmark and GO gene sets within the Molecular Signatures Database.

### RNA seq data analysis

The RNA seq reads were mapped to the mouse genome assembly mm10 using STAR version X. Repeat sequences were masked using Bowtie 2 (Langmead and Salzberg, 2012) and RepeatMasker elements (Tarailo-Graovac and Chen, 2009). Differentially expressed genes were identified using DESeq2 (Love et al., 2014) on the CD11b dataset and fdrtool (Strimmer, 2008a; Strimmer, 2008b) on the Lin-dataset. Multiple testing correction was done using the Benjamini-Hochberg method. Differentially expressed genes were considered significant if adjusted p-value < 0.1 and log2FC > 1. All data collection and parsing were completed with bash and python2.7. Statistical analyses were performed using R programming language version 3.5.1.

### Estimation of alternative splicing

Mixture of Isoforms (MISO)1 Bayesian Inference model v0.5.4 was used to quantify alternative splicing events. The MISO event database for pairwise alternative splicing events for mm10 (“exon-centric annotation”) was downloaded from hollywood.mit.edu/burgelab/miso/annotations/. After MISO quantified the percent spliced in (PSI) for each event by counting the number of reads supporting both events and the reads that are unique to each isoform, we calculated delta PSI by subtracting PSI from the WT with the I3KO sample for each alternative event. Finally, we filtered for significant and differential splicing events between wild-type and knockdown samples by requiring that delta PSI > 0.1, the Bayes factor ≥10, and the sum of exclusion and inclusion reads ≥ 10.

### Statistics

Data represent mean ±SD for continuous numerical data, unless otherwise noted in the figure legends. One-way ANOVA followed by Bonferroni’s multiple comparisons test or 2-tailed Student’s t tests were performed using GraphPad Prism software. One-way ANOVA followed by Bonferroni’s multiple comparisons test was performed in experiments with more than two groups.

### Data Sharing Statement

Raw and analyzed data have been deposited onto the NCBI Gene Expression Omnibus (GEO) repository (GSE156115).

## AUTHOR CONTRIBUTIONS

T.M.T., J.B., N.N., J.P., J.D., T.L., J.K.P., A.K.J., M.P., and J.K. performed experiments. T.M.T., J.P., and S.K. analyzed results and made the figures. O.S. provided experimental resource. T.M.T. and D.S.R. designed the research and wrote the paper. T.M.T., J.B., N.N., J.P., T.L., J.K.P., O.S., J.K., J.R.S., D.S.R. reviewed and edited the paper.

## ACKNOWLEDGEMENTS

This work was supported by the Tumor Cell Biology Training Grant NIH T32 CA009056 (T.M.T.), NIH/NIGMS R35GM130361 (J.R.S.), NIH/NCI R01CA166540 (D.S.R.), NIH/NCI R21CA197441 (D.S.R.), American Society of Hematology Bridge Grant (D.S.R.), UCLA Jonsson Comprehensive Cancer Center Seed Grant (D.S.R.) and STOPCancer/Barbara and Gary Luboff Mitzvah Fund Seed Grant (D.S.R.). Flow cytometry was performed in the Eli and Edythe Broad Center of Regenerative Medicine and Stem Cell Research UCLA Flow Cytometry Core Resource and the UCLA JCCC/CFAR Flow Cytometry Core Facility that is supported by NIH AI-28697, P30CA016042, the JCCC, the UCLA AIDS Institute, and the David Geffen School of Medicine at UCLA. The authors acknowledge the support of the Chao Family Comprehensive Cancer Center Transgenic Mouse Facility (TMF) Shared Resource, supported by the National Cancer Institute of the National Institutes of Health under award number P30CA062203. The content is solely the responsibility of the authors and does not necessarily represent the official views of the National Institutes of Health. The authors would like to thank Jon Neumann (TMF), Michael O. Alberti, and Jorge Contreras for their expertise and helpful discussions.

