## Supplemental Data for "Post-transcriptional gene regulation by the RNA binding protein IGF2BP3 is critical for MLL-AF4 mediated leukemogenesis"

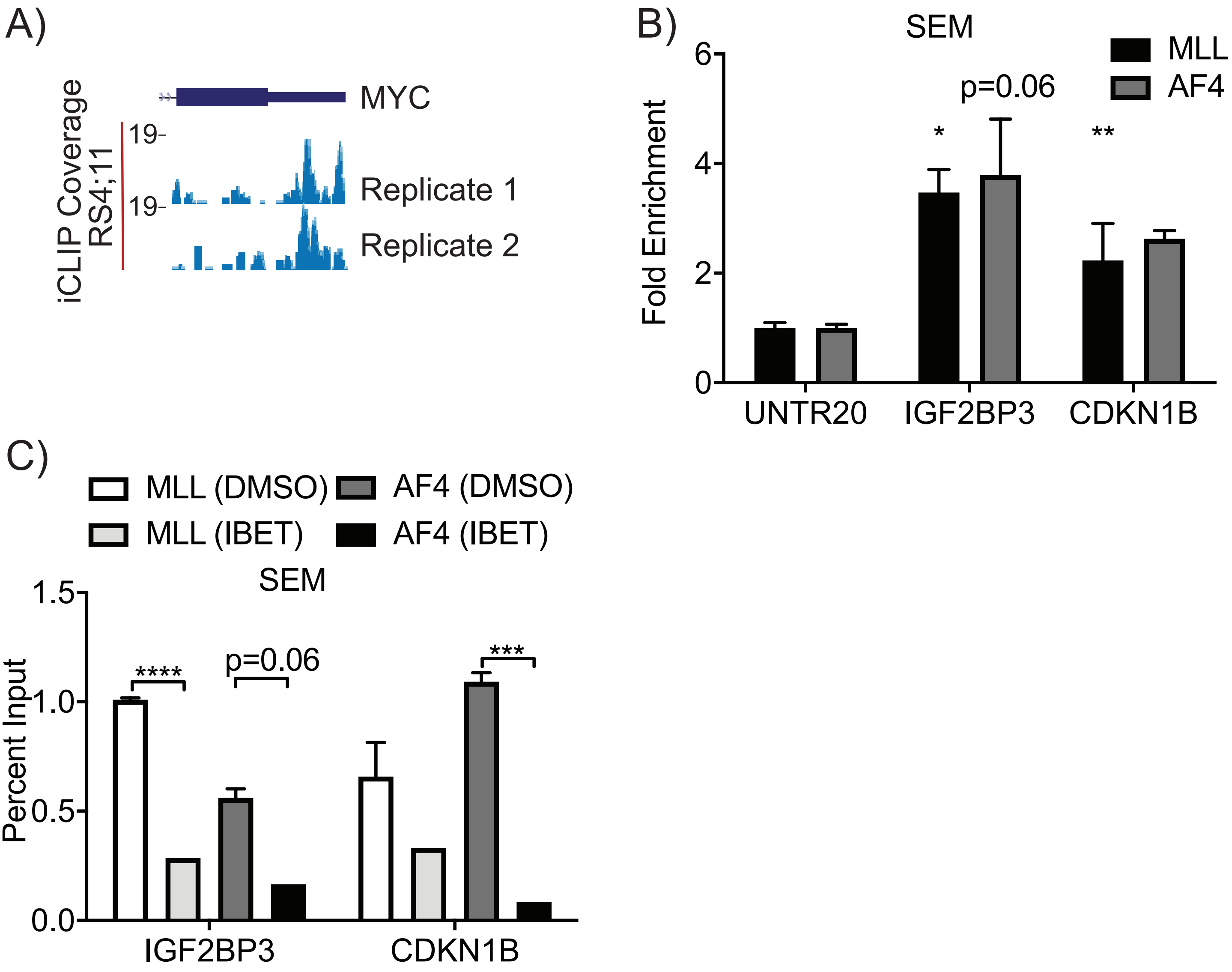

A)
MYC
iCLIP Coverage
RS4;11
19-
Replicate 1
19-
Replicate 2
B)
SEM
MLL
AF4
p=0.06
*
**
Fold Enrichment
0
2
4
6
UNTR20
IGF2BP3
CDKN1B
C)
MLL (DMSO)
AF4 (DMSO)
MLL (IBET)
AF4 (IBET)
SEM
****
p=0.06
***
Percent Input
0.0
0.5
1.0
1.5
IGF2BP3
CDKN1B

**Supplemental Figure 1, Related to Figure 1. MLL-AF4 induces IGF2BP3 and IGF2BP3 binds to MLL-AF4 transcriptional target genes.**
A) UCSC Genome Browser snapshot of RS4;11 IGF2BP3 CLIP-seq target MYC, an MLL-AF4 target gene.
B) ChIP-qPCR shows fold enrichment for IGF2BP3 and CDKN1B with MLL and AF4 IP in SEM. Normalized to UNTR20, an untranscribed region (t-test; *P < 0.05, **P < 0.01).
C) Percent input from ChIP-qPCR of SEM cells show reduced binding of MLL-AF4 to IGF2BP3 with treatment of IBET151 (t-test; *** P < 0.001, ****P < 0.0001).

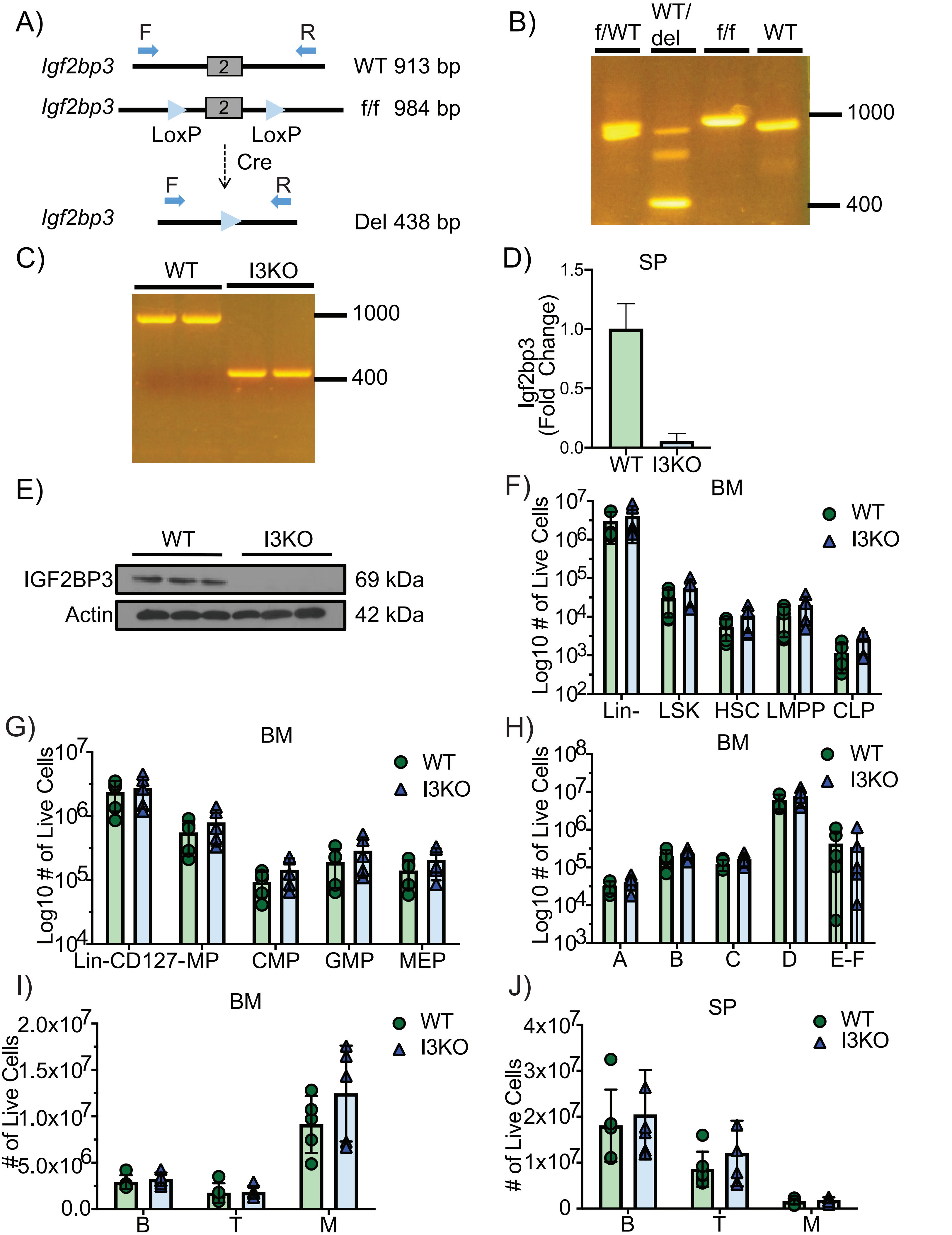
A)
F
R
Igf2bp3
2
WT 913 bp
Igf2bp3
2
f/f 984 bp
LoxP
LoxP
Cre
F
R
Igf2bp3
Del 438 bp
B)
f/WT
WT/ del
f/f
WT
1000
400
C)
WT
I3KO
1000
400
D)
SP
Igf2bp3 (Fold Change)
1.5
1.0
0.5
0.0
WT
I3KO
E)
WT
I3KO
IGF2BP3
69 kDa
Actin
42 kDa
F)
BM
WT
I3KO
Log10 # of Live Cells
Lin-
LSK
HSC
LMPP
CLP
G)
BM
WT
I3KO
Log10 # of Live Cells
Lin-CD127-MP
CMP
GMP
MEP
H)
BM
WT
I3KO
Log10 # of Live Cells
A
B
C
D
E-F
I)
BM
WT
I3KO
### of Live Cells
2.0x10^7
1.5x10^7
1.0x10^7
5.0x10^6
0.0
B
T
M
J)
SP
WT
I3KO
### of Live Cells
4x10^7
3x10^7
2x10^7
1x10^7
0
B
T
M

**Supplemental Figure 2, Related to Figure 2. I3KO mice maintain normal, steady-state hematopoiesis.**

A) Schematic of alleles in WT, *Igf2bp3*$^{f/f}$, and *Igf2bp3*$^{del/del}$ (I3KO) mice.
B) PCR genotyping showing WT, *Igf2bp3*$^{f/f}$, heterozygous I*gf2bp3*$^{f/WT}$ and heterozygous *Igf2bp3*$^{WT/del}$ alleles.
C) PCR genotyping showing WT and *Igf2bp3*$^{del/del}$ KO alleles.
D) *Igf2bp3* expression using RT-qPCR in the spleen of I3KO mice.
E) WB of IGF2BP3 expression in the spleen (tissue with the highest expression of I3) of WT and I3KO mice.
F) Quantitation of Lin-, LSKs, HSCs, LMPPs, and CLPs in the BM of WT and I3KO mice at 8 weeks (n=5 WT, n=5 I3KO; t-test; P = 0.55, 0.23, 0.16, 0.22, 0.09, respectively).
G) Quantitation of Lin-CD127-, Myeloid progenitors (MP), CMPs, GMPs, and MEPs in the BM of WT and I3KO mice at 8 weeks (n=5 WT, n=5 I3KO; t-test; P = 0.67, 0.35, 0.22, 0.34, 0.27, respectively).
H) Hardy Fractions of WT and I3KO mice at 8 weeks (n=5 WT, n=5 I3KO; t-test; P = 0.30, 0.52, 0.21, 0.44, 0.78, respectively).
I) Quantification of B cells, T cells, and Myeloid cells in the BM at 8 weeks (n=5 WT, n=5 I3KO; t-test; P = 0.51, 0.88, 0.25, respectively).
J) Quantification of B cells, T cells, and Myeloid cells in the spleen of WT and I3KO mice at 8 weeks (n=5 WT, n=5 I3KO; t-test; P = 0.70, 0.57, 0.68, respectively).

**Table S1, Related to Supplemental Figure 2. Chi-square of generation of *Igf2bp3*$^{del/del}$ (I3 KO) mice.**

| Genotype | Observed (O) | Expected (E) | O-E | $(O-E)^2$ | $(O-E)^2/E$ | $\chi^2$ | P value |
|---|---|---|---|---|---|---|---|
| WT | 20 | 14 | 6.0 | 36.0 | 2.57 | 4.39 | 0.1-0.15 |
| WT/del | 21 | 28 | -7.0 | 49.0 | 1.75 | | |
| del/del | 15 | 14 | 1.0 | 1.0 | 0.07 | | |

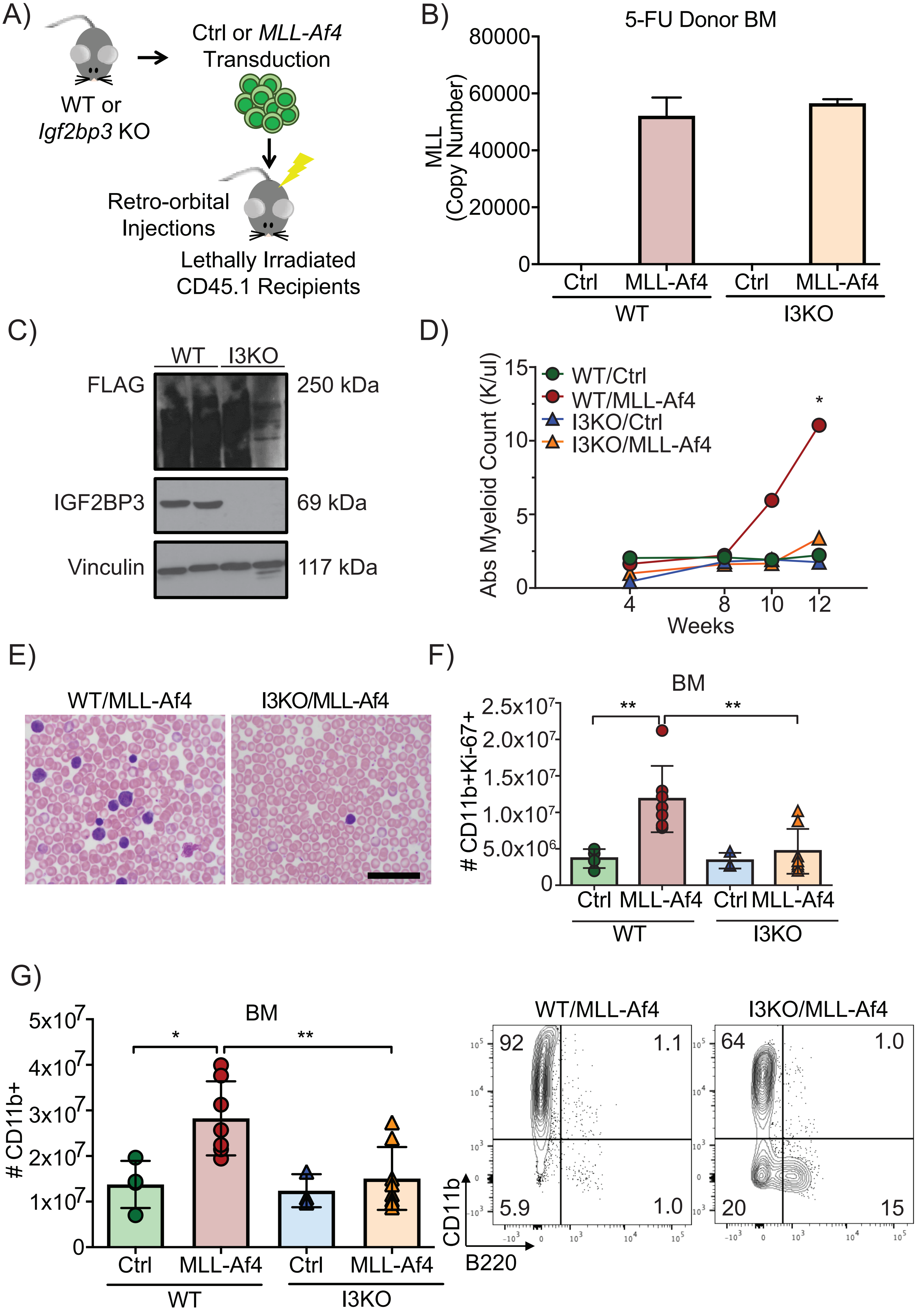
A)
WT or
Igf2bp3 KO
Ctrl or MLL-Af4
Transduction
Retro-orbital
Injections
Lethally Irradiated
CD45.1 Recipients
B)
5-FU Donor BM
MLL
(Copy Number)
80000
60000
40000
20000
0
Ctrl
MLL-Af4
Ctrl
MLL-Af4
WT
I3KO
C)
WT
I3KO
FLAG
250 kDa
IGF2BP3
69 kDa
Vinculin
117 kDa
D)
Abs Myeloid Count (K/ul)
15
10
5
0
WT/Ctrl
WT/MLL-Af4
I3KO/Ctrl
I3KO/MLL-Af4
*
4
8
10
12
Weeks
E)
WT/MLL-Af4
I3KO/MLL-Af4
F)
BM
# CD11b+Ki-67+
2.5x10^7
2.0x10^7
1.5x10^7
1.0x10^7
5.0x10^6
0
**
**
Ctrl
MLL-Af4
Ctrl
MLL-Af4
WT
I3KO
G)
BM
# CD11b+
5x10^7
4x10^7
3x10^7
2x10^7
1x10^7
0
*
**
Ctrl
MLL-Af4
Ctrl
MLL-Af4
WT
I3KO
WT/MLL-Af4
92
1.1
5.9
1.0
I3KO/MLL-Af4
64
1.0
20
15
CD11b
B220

**Supplemental Figure 3, Related to Figure 2. *Igf2bp3* deletion attenuates the disease and results in the reduction of leukemic cells.**

A) Schematic of BMT.

B) MLL-Af4 copy number in 5-FU treated bone marrow from CD45.2 donor mice.

C) Western blot of BM from mice transplanted with MLL-Af4 transduced WT or I3KO donor HSPCs with antibodies to FLAG, IGF2BP3 and Vinculin.

D) Time course of Absolute Myeloid Counts in the PB of mice transplanted with control (Ctrl) or MLL-Af4 transduced HSPCs from WT or I3KO mice (t-test; *$P < 0.05$).

E) Wright staining of PB smears from WT/MLL-Af4 and I3KO/MLL-Af4 mouse. Scale bar, 40 microns

F) Quantitation of CD11b+Ki67+ cells in the BM at 14 weeks post-transplantation (n= 4 Ctrl, n=8 MLL-Af4; one-way ANOVA followed by Bonferroni's multiple comparisons test; **$P < 0.01$).

G) (Left) Number of CD11b+ in the BM of recipient mice that received Ctrl or MLL-Af4 transduced HSPCs from WT or I3KO mice at 14 weeks (n=4 Ctrl, n=8 MLL-Af4; one-way ANOVA followed by Bonferroni's multiple comparisons test; *$P < 0.05$, **$P < 0.01$). (Right) Corresponding representative FACS plots showing CD11b+ and B220+ cells in the BM.

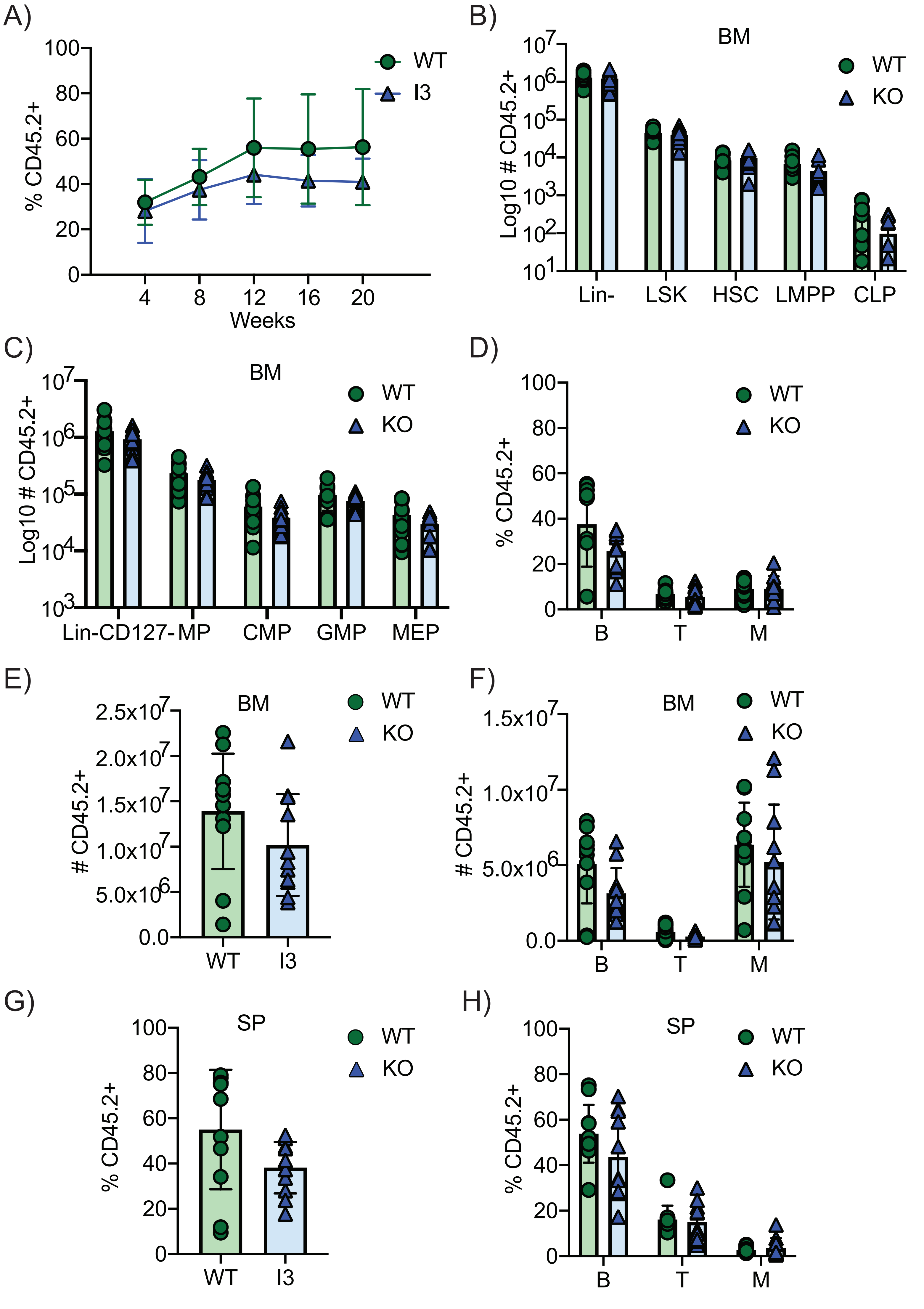
A)
% CD45.2+
Weeks
WT
I3
B)
BM
Log10 # CD45.2+
WT
KO
Lin-
LSK
HSC
LMPP
CLP
C)
BM
Log10 # CD45.2+
WT
KO
Lin-CD127-MP
CMP
GMP
MEP
D)
% CD45.2+
WT
KO
B
T
M
E)
BM
# CD45.2+
WT
KO
WT
I3
F)
BM
# CD45.2+
WT
KO
B
T
M
G)
SP
% CD45.2+
WT
KO
WT
I3
H)
SP
% CD45.2+
WT
KO
B
T
M

**Supplemental Figure 4, Related to Figure 3-4. I3KO mice show normal hematopoietic stem cell function and ability to reconstitute mice.**

A) Percentage of CD45.2+ engraftment in the PB of recipient mice transplanted with either WT or I3KO donor BM in competitive repopulation BMT (n=11 WT, n=11 I3KO; t-test; P = 0.47, 0.31, 0.13, 0.1, 0.08, respectively).

B) Quantitation of Lin-, LSKs, HSCs, LMPPs, and CLPs in the BM of competitive repopulation BMT mice at 24 weeks post-transplantation (n=11 WT, n=11 I3KO; t-test; P = 0.92, 0.51, 0.42, 0.13,0. 051, respectively).

C) Quantitation of Lin-CD127-, Myeloid progenitors (MP), CMPs, GMPs, and MEPs in the BM of competitive repopulation BMT at 24 weeks (n=11 WT, n=11 I3KO; t-test; P = 0.17, 0.18, 0.09, 0.18, 0.17, respectively).

D) Percentage of CD45.2+ B cells (B), T cells (T), Myeloid cells (M) in the PB of competitive repopulation BMT mice at 24 weeks (n=11 WT, n=11 I3KO; t-test; P = 0.07, 0.34, 0.95, respectively).

E) Quantification of CD45.2+ cells in the BM of competitive repopulation BMT mice at 24 weeks (n=11 WT, n=11 I3KO; t-test; P =0.16).

F) Quantification of CD45.2+ B cells (B), T cells (T), Myeloid cells (M) in the BM of competitive repopulation BMT mice at 24 weeks (n=11 WT, n=11 I3KO; t-test; P = 0.051, 0.06, 0.43, respectively).

G) Percentage of CD45.2+ cells in the SP of competitive repopulation BMT mice at 24 weeks (n=11 WT, n=11 I3KO; t-test; P =0.07).

H) Percentage of CD45.2+ B cells (B), T cells (T), Myeloid cells (M) in the SP of competitive repopulation BMT mice at 24 weeks (n=11 WT, n=11 I3KO; t-test; P = 0.14, 0.72, 0.44, respectively).

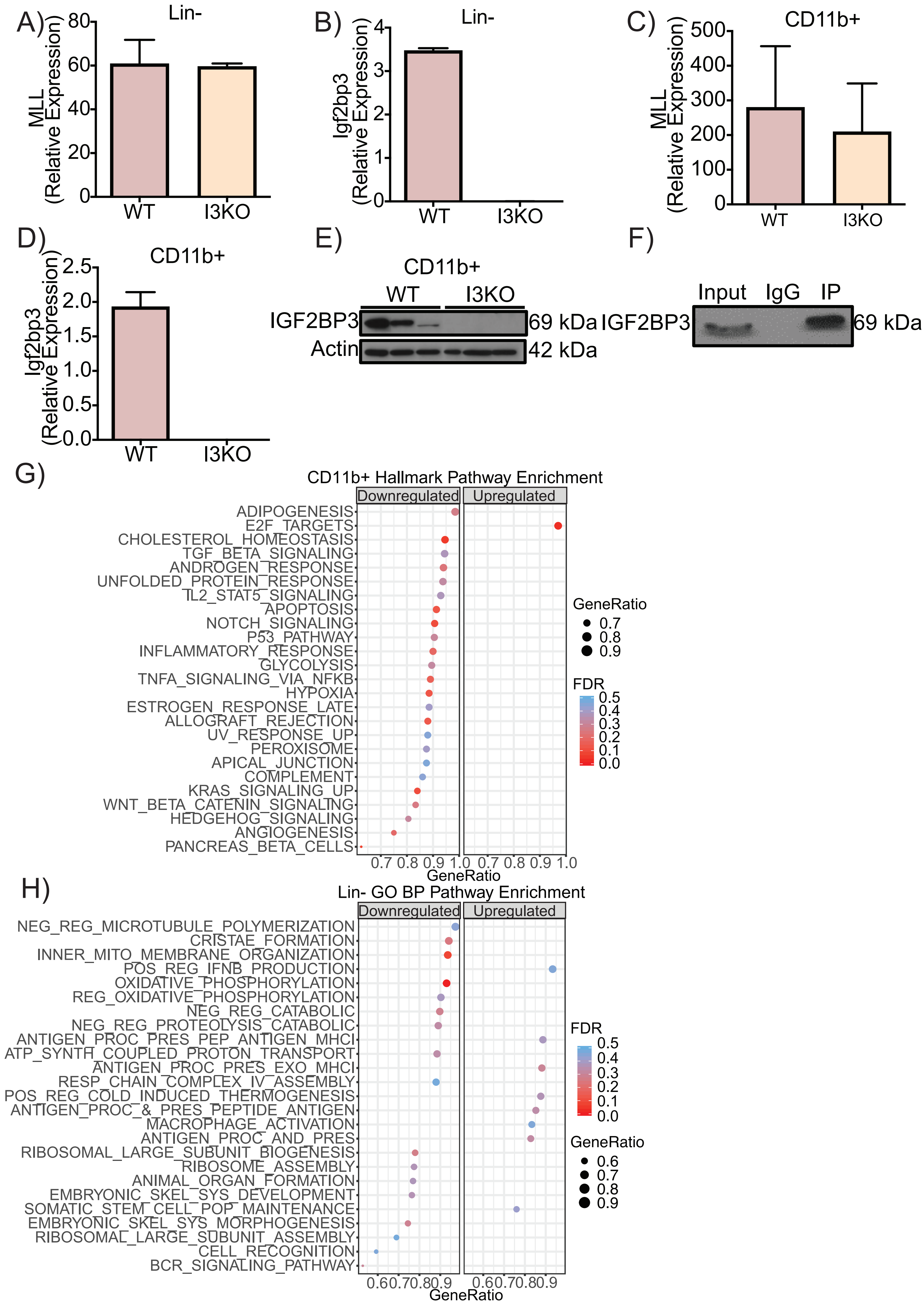
A)
Lin-
MLL (Relative Expression)
0
20
40
60
80
WT
I3KO
B)
Lin-
Igf2bp3 (Relative Expression)
0
1
2
3
4
WT
I3KO
C)
CD11b+
MLL (Relative Expression)
0
100
200
300
400
500
WT
I3KO
D)
CD11b+
Igf2bp3 (Relative Expression)
0.0
0.5
1.0
1.5
2.0
2.5
WT
I3KO
E)
CD11b+
WT
I3KO
IGF2BP3
Actin
69 kDa
42 kDa
F)
Input
IgG
IP
IGF2BP3
69 kDa
G)
CD11b+ Hallmark Pathway Enrichment
Downregulated
Upregulated
ADIPOGENESIS
E2F_TARGETS
CHOLESTEROL_HOMEOSTASIS
TGF_BETA_SIGNALING
ANDROGEN_RESPONSE
UNFOLDED_PROTEIN_RESPONSE
IL2_STAT5_SIGNALING
APOPTOSIS
NOTCH_SIGNALING
P53_PATHWAY
INFLAMMATORY_RESPONSE
GLYCOLYSIS
TNFA_SIGNALING_VIA_NFKB
HYPOXIA
ESTROGEN_RESPONSE_LATE
ALLOGRAFT_REJECTION
UV_RESPONSE_UP
PEROXISOME
APICAL_JUNCTION
COMPLEMENT
KRAS_SIGNALING_UP
WNT_BETA_CATENIN_SIGNALING
HEDGEHOG_SIGNALING
ANGIOGENESIS
PANCREAS_BETA_CELLS
0.7 0.8 0.9 1.0 0.7 0.8 0.9 1.0
GeneRatio
GeneRatio
0.7
0.8
0.9
FDR
0.5
0.4
0.3
0.2
0.1
0.0
H)
Lin- GO BP Pathway Enrichment
Downregulated
Upregulated
NEG_REG_MICROTUBULE_POLYMERIZATION
CRISTAE_FORMATION
INNER_MITO_MEMBRANE_ORGANIZATION
POS_REG_IFNB_PRODUCTION
OXIDATIVE_PHOSPHORYLATION
REG_OXIDATIVE_PHOSPHORYLATION
NEG_REG_CATABOLIC
NEG_REG_PROTEOLYSIS_CATABOLIC
ANTIGEN_PROC_PRES_PEP_ANTIGEN_MHCI
ATP_SYNTH_COUPLED_PROTON_TRANSPORT
ANTIGEN_PROC_PRES_EXO_MHCI
RESP_CHAIN_COMPLEX_IV_ASSEMBLY
POS_REG_COLD_INDUCED_THERMOGENESIS
ANTIGEN_PROC_&_PRES_PEPTIDE_ANTIGEN
MACROPHAGE_ACTIVATION
ANTIGEN_PROC_AND_PRES
RIBOSOMAL_LARGE_SUBUNIT_BIOGENESIS
RIBOSOME_ASSEMBLY
ANIMAL_ORGAN_FORMATION
EMBRYONIC_SKEL_SYS_DEVELOPMENT
SOMATIC_STEM_CELL_POP_MAINTENANCE
EMBRYONIC_SKEL_SYS_MORPHOGENESIS
RIBOSOMAL_LARGE_SUBUNIT_ASSEMBLY
CELL_RECOGNITION
BCR_SIGNALING_PATHWAY
0.6 0.7 0.8 0.9 0.6 0.7 0.8 0.9
GeneRatio
FDR
0.5
0.4
0.3
0.2
0.1
0.0
GeneRatio
0.6
0.7
0.8
0.9

**Supplemental Figure 5, Related to Figure 3 and 5. *Igf2bp3* is upregulated in MLL-Af4 leukemia cells and enhances leukemogenesis by targeting transcripts within Ras signaling pathways.**

A) Relative expression of MLL by RT-qPCR in WT/MLL-Af4 and I3KO/MLL-Af4 immortalized Lin- cells.

B) Relative expression of *Igf2bp3* by RT-qPCR in WT/MLL-Af4 and I3KO/ MLL-Af4 immortalized Lin- cells.

C) Relative expression of *MLL* by RT-qPCR in WT/MLL-Af4 and I3KO/MLL-Af4 leukemic CD11b+ splenic cells.

D) Relative expression of *Igf2bp3* by RT-qPCR in WT/MLL-Af4 and I3KO/MLL-Af4 leukemic CD11b+ splenic cells.

E) Expression of IGF2BP3 of in WT/MLL-Af4 and I3KO/MLL-Af4 leukemic CD11b+ splenic cells at the protein level.

F) Confirmation Western blot of protein samples from IGF2BP3 RIP. Input refers to 70Z/3 cell lysate used for immunoprecipitation. RIP is RNA immunoprecipitation from control (mouse IgG) or α-IGF2BP3 antibody used in eCLIP (MBL).

G) Hallmark Pathway enrichment using Gene Set Enrichment Analysis on DEseq analyzed RNA-seq samples from WT/MLL-Af4 and I3KO/MLL-Af4 CD11b+ cells.

H) Gene Ontology Biological Processes Pathway enrichment using Gene Set Enrichment Analysis on DEseq analyzed RNA-seq samples from WT/MLL-Af4 and I3KO/MLL-Af4 Lin- cells.

**Table S2, Related to Figure 5-6. IGF2BP3-dependent regulatory targets in MLL-Af4 Lin- cells.** Provided as an Excel file.

**Table S3, Related to Figure 5-6. IGF2BP3-dependent regulatory targets in MLL-Af4 CD11b+ bulk leukemia cells.** Provided as an Excel file.

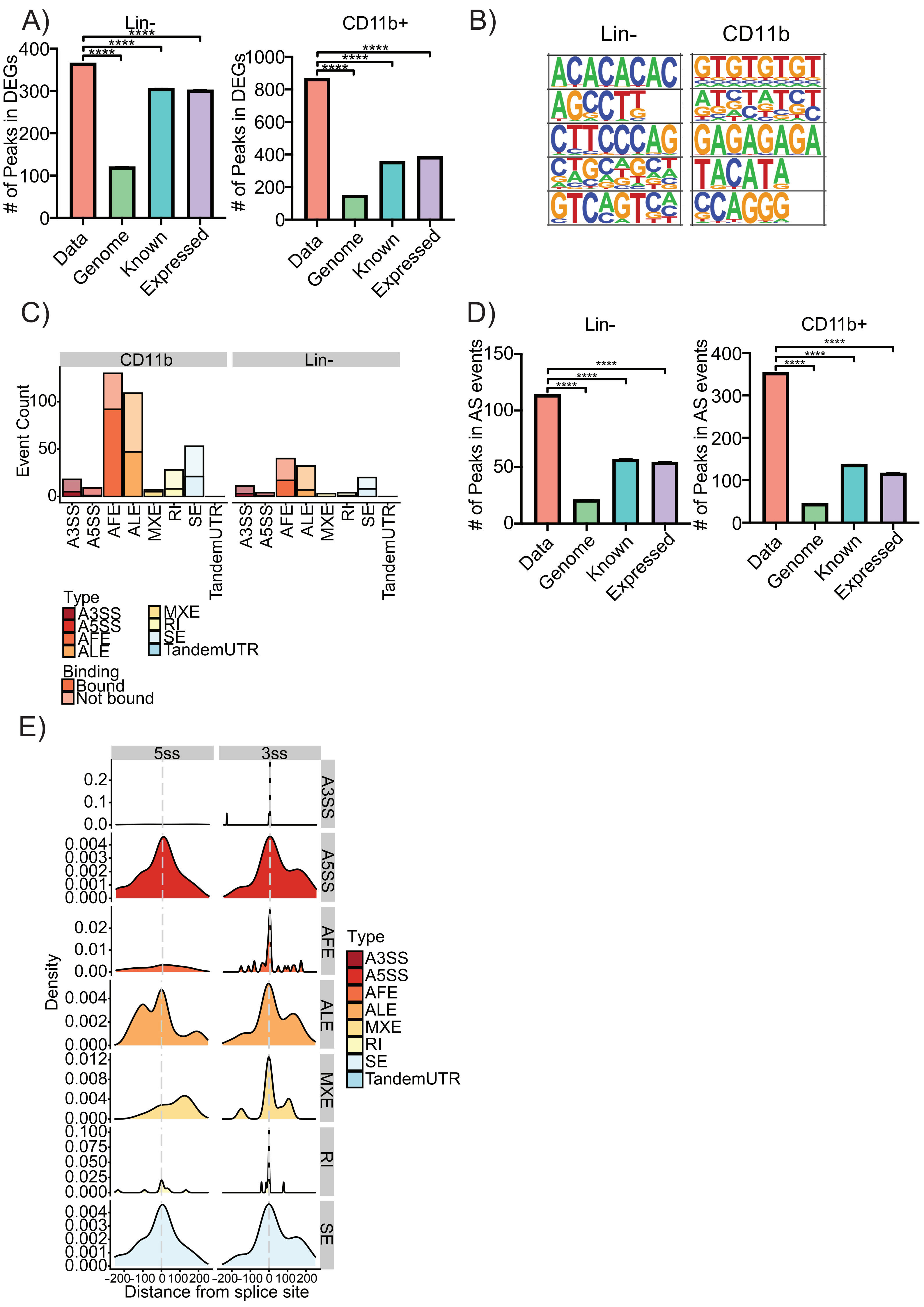

A)
Lin-
CD11b+
### of Peaks in DEGs
Data
Genome
Known
Expressed
B)
Lin-
CD11b
C)
CD11b
Lin-
Event Count
A3SS
A5SS
AFE
ALE
MXE
RI
SE
TandemUTR
Type
Binding
Bound
Not bound
D)
Lin-
CD11b+
### of Peaks in AS events
E)
5ss
3ss
Density
Distance from splice site

**Supplemental Figure 6, Related to Figure 6**. ***Igf2bp3* targets specific mRNA regulons through multiple post-transcriptional mechanisms.**
A) Number of predicted eCLIP binding sites within the differentially expressed genes in the Lin- (left) and CD11b+ (right) dataset, the whole genome, known genes, and expressed genes by simulating distribution of binding sites over respective background (data represent mean±SEM; one-sample t-test; ****$P < 2.2x10^{-16}$).
B) Homer motif analysis showing top five 6-8-mer motifs.
C) Event counts for different types of alternative splicing patterns in CD11b and Lin- datasets (A3SS, Alternative 3' splice sites; A5SS, Alternative 5' splice sites; AFE, Alternative first exons; ALE, Alternative last exons; MXE, Mutually exclusive exons; RI, Retained introns; SE, Skipped exons; Bound, IGF2BP3 eCLIP target).
D) Number of predicted eCLIP binding sites within the dynamic alternative splicing events in the Lin- (left) and CD11b+ dataset (right), the whole genome, known genes, and expressed genes by simulating distribution of binding sites over respective background (data represent mean±SEM; one-sample t-test; ****$P < 2.2x10^{-16}$).
E) Histogram showing alternative event density and distance from 5' (5ss) and 3' (3ss) splice sites for each alternative splicing pattern (A3SS, Alternative 3' splice sites; A5SS, Alternative 5' splice sites; AFE, Alternative first exons; ALE, Alternative last exons; MXE, Mutually exclusive exons; RI, Retained introns; SE, Skipped exons).

#### Table S4, Related to Methods. Antibodies and oligonucleotides used in experiments.

| REAGENT | VENDOR/REFERENCE | CATALOG NUMBER |
|---|---|---|
| Antibodies | | |
| MLL rabbit polyclonal | Bethyl Lab | A300-086A |
| AF4 rabbit polyclonal | Abcam | ab31812 |
| RNA Polymerase mouse monoclonal | EMD Millipore | 05-623B |
| Normal Mouse IgG | EMD Millipore | 12-371B |
| FLAG mouse monoclonal | Sigma-Aldrich | F1804 |
| IGF2BP3 rabbit polyclonal | MBL | RN009P |
| Vinculin mouse monoclonal | Santa Cruz Biotechnology | sc-73614 |
| β-actin mouse monoclonal | Sigma-Aldrich | A1978 |
| HOXA9 goat polyclonal | Santa Cruz Biotechnology | sc-17155 |
| CD3e-PE | Biolegend | 100308 |
| CD11b-PE-Cy7 | Biolegend | 101216 |
| B220-PercP-Cy5.5 | Biolegend | 103236 |
| Gr-1-Pacific Blue | Biolegend | 108430 |
| CD45.1-APC-Cy7 | Biolegend | 110716 |
| CD45.2-APC | Biolegend | 109813 |
| c-Kit/CD117-Alexa Fluor 700 | Thermo Fisher Scientific | 56-1172-80 |
| Ki-67-PE | Biolegend | 652403 |
| c-Kit/CD117-APC-Cy7 | Biolegend | 105826 |
| Sca1-PerCP-Cy5.5 | Biolegend | 108124 |
| CD135- APC | Thermo Fisher Scientific | 17-1351-82 |
| CD127-PE-Cy7 | Biolegend | 135014 |
| CD150-PE | Biolegend | 115903 |
| Biotin CD4 | Thermo Fisher Scientific | 13-0041-82 |
| Biotin CD8 | Biolegend | 100704 |
| Biotin B220 | Biolegend | 103204 |
| Biotin NK1.1 | Biolegend | 108704 |
| Biotin IgM | Thermo Fisher Scientific | 13-5790-85 |
| Biotin Gr-1 | Biolegend | 108404 |
| Biotin Ter119 | Biolegend | 116204 |
| Biotin TCR beta | Biolegend | 109204 |
| Biotin TCR gamma-delta | Biolegend | 118103 |
| CD45.2-BV605 | Biolegend | 109841 |
| Streptavidin-Pacific Blue | Thermo Fisher Scientific | 48-4317-82 |
| CD16/32-PE-Cy7 | Biolegend | 101318 |
| CD34-Alexa Fluor 700 | Thermo Fisher Scientific | 56-0341-82 |
| CD127-APC | Biolegend | 135012 |
| IgM-PE | SoutherBiotech | 1020-09S |
| CD43- APC | BD Biosciences | 560663 |
| CD24-PE-Cy7 | Thermo Fisher Scientific | 25-0242-80 |
| Ly51-Biotin | Biolegend | 108303 |

| Oligonucleotides | | |
|---|---|---|
| IGF2BP3-ChIP F: GACCACGAACGGGAGAACTG R: TCAATTCAGACGTGGTGCGG | Lin et al., 2016. | N/A |
| CDKN1B-ChIP F: TCTTCTTCGTCAGCCTCCCTTC R: TCGCAGAGCCGTGAGCAAGC | Wilkinson et al., 2013. | N/A |
| MLL F: GAAACCTACCCCATCAGCAA R: GACCTGCTTGCTTGACTTCC | This paper | N/A |
| Igf2bp3 F: CCTGGTGAAGACGGGCTAC R: TCAACTTCCATCGGTTTCCCA | This paper | N/A |
| Hoxa7 F: ATGTGAACGCGCTTTTTAGC R: ATTGTATAAGCCCGGCACAG | This paper | N/A |
| Hoxa9 F: AAAACACCAGACGCTGGAAC R: TCTTTTGCTCGGTCCTTGTT | This paper | N/A |
| Hoxa10 F: GAAGAAACGCTGCCCTTACA R: GATTCGGTTTTCTCGGTTCA | This paper | N/A |
| Ccnd1 F: GCGTACCCTGACACCAATCT R: CTCTTCGCACTTCTGCTCCT | This paper | N/A |
| Maf F: GAGGAGGTGATCCGACTGAA R: TCTCCTGCTTGAGGTGGTCT | This paper | N/A |
| Itga6 F: TGAAGATGGGCCCTATGAAG R: CTCTTGGAGCACCAGACACA | This paper | N/A |
| Igf2bp3 sgRNA F:CACCGAGCTTGGTCCTTACTGGAAT R:AAACATTCCAGTAAGGACCAAGCTC | Palanichamy et al., 2016. | N/A |
| Non-targeting sgRNA F:TTTGCGAGGTATTCGGCTCCGCG R: AAACCGCGGAGCCGAATACCTCG | Sanjana et al., 2014. | N/A |
| L32 F: AAGCGAAACTGGCGGAAAC R: TAACCGATGTTGGGCATCAG | Palanichamy et al., 2016. | N/A |
| UNTR20 F: ATCACACTGCAAAAATCCAGAA R: TCACTTCTTTAACTGGCCTTGA | Kaukonen et al., 2016. | N/A |

#### SUPPLEMENTAL METHODS

*ChIP-PCR reagents.* Chromatin immunoprecipitation was performed using the EZ-ChIP kit (EMD Millipore 17-371). Antibodies used were MLL rabbit polyclonal (Bethyl Lab A300-086A), AF4 rabbit polyclonal (Abcam ab31812), RNA Polymerase mouse monoclonal (EMD Millipore 05-623B), and Normal Mouse IgG (EMD Millipore 12-371B). SEM cells were treated with 1μM of I-BET151 (Sigma-Aldrich SML0666).

*qPCR reagents.* RNA from cell lines or primary mouse cells was collected using QIAzol Lysis Reagent and purified using the miRNeasy Mini Kit (Qiagen 217004). RNA was reverse transcribed using qScript (Quanta BioSciences). qPCR was performed with the StepOne Plus Real-Time PCR System (Applied Biosystems) using PerfeCTa SYBR Green FastMix reagent (Quanta BioSciences).

*Western blot reagents.* Cells were lysed in RIPA buffer (Boston BioProducts) supplemented with Halt Protease and Phosphatase Inhibitor Cocktail (Thermo Scientific). Equal amounts of protein lysate (as quantified by using bicinchoninic acid protein assay, BCA [Thermo Scientific]) were electrophoresed on a 5%–12% SDS-PAGE and electroblotted onto a nitrocellulose membrane. Antibodies used were FLAG mouse monoclonal (Sigma-Aldrich F1804), IGF2BP3 rabbit polyclonal (MBL RN009P), MLL rabbit polyclonal (Bethyl Lab A300-086A), HOXA9 goat polyclonal (Santa Cruz Biotechnology Inc. sc-17155), Vinculin mouse monoclonal (Santa Cruz Biotechnology Inc. sc-73614) and β-actin mouse monoclonal (Sigma-Aldrich A1978). Secondary HRP-conjugated antibodies (Santa Cruz Biotechnology Inc.) and SuperSignal West Pico Plus kit (Pierce Biotechnology) were used for enhanced chemiluminescence-based detection. Quantification was performed using ImageJ analysis.

*Luciferase assays.* The 950bp promoter region upstream of the TSS of IGF2BP3 was cloned into the multiple cloning region between NheI and HindIII restriction enzyme sites of the pGL4.11 vector (Promega). 293T cells were cotransfected with the pGL4.11, pGL.4.75, IGF2BP3 promoter containing reporter vector along with the empty vector or PIDE-MLL-AF4 overexpression vector ratios (0:400, 375:25, 350:50, 300:100, 200:200, and 0:400 ng). Cotransfections were performed with BioT (Bioland) as per the manufacturer's instructions. Cells were lysed after 24 hours, substrate was added, and luminescence was measured on a GloMax-Multi Jr (Promega) utilizing the Dual-Luciferase Reporter Assay System (Promega E1910). The ratio of firefly to Renilla luciferase activity was calculated for all samples.

*Additional plasmids, cell culture, and spin infection details.* The PIDE-MLL-AF4 plasmid used in the luciferase assay was kindly given by Rolf Marschalek (Goethe-University of Frankfurt, Frankfurt, Germany) through MTA[1]. 70Z/3 cells (ATCC TIB-158) were spin-infected at 30°C for 90 minutes in the presence of polybrene. Lin- cells were obtained by harvesting the BM from C57BL/6J (Jackson Laboratory 000664) or I3KO mice, lysing in red blood cell lysis buffer, and staining with a cocktail of biotinylated antibodies for lineage depletion by MACS (Miltenyi). Cas9 Lin- cells were obtained by harvesting the BM from B6J.129(Cg)-Gt(ROSA)26Sor$^{tm1.1(CAG\text{-}cas9^{*},\text{-}EGFP)Fezh}$/J mice (Jackson Laboratory 026179), lysing in red blood cell lysis buffer, and staining with a cocktail of biotinylated antibodies for lineage depletion by MACS (Miltenyi). After transduction and selection as described above, Cas9 Lin- cells were spin-infected with MSCV-hU6-NT/I3sgRNA-EFS-mCherry virus at 30°C for 90 minutes in the presence of polybrene. CD11b+ cells were obtained by harvesting the splenic tumors from the WT/MLL-Af4 or I3KO/MLL-Af4 mice, lysing in red blood cell lysis buffer, and staining with a CD11b biotinylated antibody (BioLegend 101204) for positive selection by MACS (Miltenyi). The list of antibodies used is provided in the Table S1. The human B-ALL cell lines RS4;11 (MLL-AF4 translocated; ATCC CRL-1873), SEM (MLL-AF4 translocated; DSMZ ACC 546), murine pre-B leukemic cell line 70Z/3 (ATCC TIB- 158), and HEK 293T cell line (ATCC CRL-11268) were grown in their corresponding media at 37°C in a 5% CO2 incubator. Lentiviruses and retroviruses were generated as previously described[2,3].

*Additional BM transplant and competitive repopulation assay details.* 5-FU enriched BM was harvested and spin-infected from 8-week-old CD45.2+ donor C57BL/6J (Jackson Laboratory 000664) or I3KO female mice as previously described[2]. 8-week-old CD45.1+ recipient B6.SJL-Ptprc-Pep3/BoyJ (Jackson Laboratory 002014) female mice were lethally irradiated and injected with donor BM. Eight mice were used per group. For competitive repopulation experiments, 8-week-old CD45.2+ donor C57BL/6J or I3KO female mice and 8-week- old CD45.1+ donor B6.SJL-Ptprc-Pep3/BoyJ female mice were harvested for BM. CD45.1+ and CD45.2+ BM cells were mixed in a ratio of 1:1 and injected into lethally irradiated 8-week-old CD45.1+ recipient B6.SJL-Ptprc-Pep3/BoyJ female mice. Mice were bled at 4, 8, 10, 12, 14, 16, and 20 weeks after BM injection for analysis of the peripheral blood. For serial secondary transplantation assays, WT/MLL-Af4 or I3KO/MLL-Af4 mice that succumbed to leukemia at 10-14 weeks post-transplantation were euthanized and the BM was collected. For each group, $10^6$, $10^5$, or $10^4$ donor BM cells were injected per mouse into 8-week-old immunocompetent CD45.1+ recipient B6.SJL-Ptprc-Pep3/BoyJ female mice. Mice were bled at 2 and 4 weeks after BM injection for analysis of the peripheral blood. All mice were purchased from the Jackson Laboratory and housed under pathogen-free conditions at UCLA.

*Colony formation assays.* WT and I3KO Lin- cells were sorted as described previously and 100,000 cells were plated on M3434 (MethoCult) methylcellulose medium containing SCF, IL-3, IL-6, and EPO. Following G418 selection, 25,000 MLL-Af4 expressing WT or I3KO Lin- cells were serially plated on M3434 methylcellulose medium. Colonies were counted at 7 and 14 days after plating.

*Intracellular flow cytometry.* For intracellular staining, after initial staining with surface marker antibodies and fixation with IC Fixation Buffer (Thermo Scientific) cells were incubated with antibodies against intracellular antigens Ki67 (BioLegend 652403) with 1X Permeabilization Buffer (Thermo Fisher Scientific). After 30 minutes of staining at RT, cells were washed twice with Permeabilization Buffer and fixed with 1% PFA. Flow cytometry was performed at the Eli and Edythe Broad Center of Regenerative Medicine and Stem Cell Research UCLA Flow Cytometry Core and the UCLA JCCC Flow Core on a BD FACS LSRII instrument. Analysis was performed using FlowJo software.

*eCLIP Library Preparation.* Following immunoprecipitation, two percent of each sample was then removed for paired input libraries. Immunoprecipitated IGF2BP3 and crosslinked RNA were run on and excised from an acrylamide gel. Library preparation was performed as outlined in the Eclipse BioInnovations Inc eCLIP kit (Eclipse BioInnovations ECEK-0001) and final libraries were excised from an acrylamide gel.

*eCLIP statistical details.* Peaks were removed if they overlapped a minimal amount of the peak length (0.0001). Peaks were further filtered for a p-value < 0.05. The motif analysis was conducted for 4-6 and 6-8bp long motifs over the HOMER standard shuffled background. Distances from peaks to splice sites were determined using bedtools closest.

*RNA seq library preparation.* The Agilent 2100 Bioanalyzer using RNA pico chip was utilized to quality check total RNA. The Universal plus mRNA-Seq Kit (NuGEN Technologies 0520-24) was used to generate strand-specific RNA-seq libraries. Samples were subjected to poly(A) RNA selection, RNA fragmentation and double-stranded cDNA generation using a mixture of random and oligo(dT) priming. The cDNA samples were then subjected to end repair to generate blunt ends, adapter ligation, strand selection, and PCR amplification to generate the final library. To mulitplex samples in one sequencing lane, different index adapters were used.

*Survival data analysis.* Survival data was computed using the Kaplan-Meier method and survival curves were compared using a Log-rank test on GraphPad Prism software.


1. Bursen A, Schwabe K, Rüster B, et al. The AF4·MLL fusion protein is capable of inducing ALL in mice without requirement of MLL·AF4. *Blood.* 2010;115(17):3570-3579.
2. O'Connell RM, Balazs AB, Rao DS, Kivork C, Yang L, Baltimore D. Lentiviral Vector Delivery of Human Interleukin-7 (hIL-7) to Human Immune System (HIS) Mice Expands T Lymphocyte Populations. *PLOS ONE.* 2010;5(8):e12009.
3. Rao DS, O'Connell RM, Chaudhuri AA, Garcia-Flores Y, Geiger TL, Baltimore D. MicroRNA-34a Perturbs B Lymphocyte Development by Repressing the Forkhead Box Transcription Factor Foxp1. *Immunity.* 2010;33(1):48-59.
